# An anatomically accurate and personalizable head injury model: Significance of brain and white matter tract morphological variability on strain

**DOI:** 10.1101/2020.05.20.105635

**Authors:** Xiaogai Li, Zhou Zhou, Svein Kleiven

**Affiliations:** Division of Neuronic Engineering, Department of Biomedical Engineering and Health Systems, KTH Royal Institute of Technology, Huddinge 141 52, Sweden

**Keywords:** Traumatic brain injury, subject-specific head model, Demons and Dramms image registration, mesh morphing, finite element analysis

## Abstract

Finite element head (FE) models are important numerical tools to study head injuries and develop protection systems. The generation of anatomically accurate and subject-specific head models with conforming hexahedral meshes remains a significant challenge. The focus of this study is to present two developmental work: First, an anatomically detailed FE head model with conforming hexahedral meshes that has smooth interfaces between the brain and the cerebrospinal fluid, embedded with white matter (WM) fiber tracts; Second, a morphing approach for subject-specific head model generation via a new hierarchical image registration pipeline integrating Demons and Dramms deformable registration algorithms. The performance of the head model is evaluated by comparing model predictions with experimental data of brain-skull relative motion, brain strain, and intracranial pressure. To demonstrate the applicability of the head model and the pipeline, six subject-specific head models of largely varying intracranial volume and shape are generated, incorporated with subject-specific WM fiber tracts. DICE similarity coefficients for cranial, brain mask, local brain regions, and lateral ventricles are calculated to evaluate personalization accuracy, demonstrating the efficiency of the pipeline in generating detailed subject-specific head models achieving satisfactory element quality without further mesh repairing. The six head models are then subjected to the same concussive loading to study sensitivity of brain strain to inter-subject variability of the brain and WM fiber morphology. The simulation results show significant differences in maximum principal strain (MPS) and axonal strain (MAS) in local brain regions (one-way ANOVA test, p<0.001), as well as their locations also vary among the subjects, demonstrating the need to further investigate the significance of subject-specific models. The techniques developed in this study may contribute to better evaluation of individual brain injury and development of individualized head protection systems in the future. This study also contains general aspects the research community may find useful: on the use of experimental brain strain close to or at injury level for head model validation; the hierarchical image registration pipeline can be used to morph other head models, such as smoothed-voxel models.

## 1 Introduction

Traumatic brain injury (TBI) is a leading cause of injury-related death and disability, with a devastating impact on the patients and their families (Maas et al. 2017). TBI causes a substantial threat to global public health and enormous economic burden for the society, with an estimated number of 50–60 million new TBI cases occur annually (Feigin et al. 2013). TBI influences all age groups, including children, adolescents, and the elderly, which can happen due to traffic accidents, sports injuries, and falls. Studies also suggest that TBI might represent an important modifiable risk factor for epilepsy, stroke, and late-life neurodegenerative diseases such as dementia and Parkinson’s disease (Maas et al. 2017). Biomechanical studies, including experimental and computational studies, have long been carried out to understand brain injury mechanisms due to external mechanical input (Meaney et al. 2014). In particular, finite element (FE) head models have emerged as valuable numerical tools to study head injuries and aid the development of protection systems (Giudice et al. 2019; Horstemeyer et al. 2019; Madhukar and Ostoja-Starzewski 2019).

Human head models with varying levels of anatomical accuracy and modeling complexity have been developed during the recent decades, e.g., WSUBIM (Ruan et al. 1994; Zhang et al. 2001), SUFEHM (earlier called ULP) (Kang et al. 1997; Sahoo et al. 2014), KTH head model (Kleiven and von Holst 2002; Kleiven 2007; Giordano and Kleiven 2014b; Zhou et al. 2019a), UCDBTM (Horgan and Gilchrist 2003; Trotta et al. 2020), SIMon (Takhounts et al. 2003; Takhounts et al. 2008), THUMS (Kimpara et al. 2006; Atsumi et al. 2016), GHBMC (Mao et al. 2013; Wu et al. 2019), WHIM (earlier called DHIM) (Ji et al. 2015; Zhao and Ji 2018; Zhao and Ji 2020). Continued efforts on model enhancement, including material model improvement, incorporating diffusion tensor imaging (DTI), brain-skull interface improvement, as well as mesh refinement, have led to updated versions compared to the original. Especially recent efforts on mesh refinement led to average brain element size of about 1.8 mm in the WHIM V1.5 model (Zhao and Ji 2019; Zhao and Ji 2020), while the element sizes in GHBMC (Mao et al. 2013a) and the refined THUMS (Atsumi et al. 2016) are also on the order of millimeter, being 2 mm, and 1.2∼5 mm, respectively. However, the brains in the above-mentioned models are simplified by smoothing out sulci and gyri, accompanied by a homogenous layer of outer cerebrospinal fluid (CSF). Further, the brain ventricles often lack anatomical details and also have jagged interface connecting with neighboring brain elements; some models or earlier versions have no ventricles. Mesh simplification as such is partially attributed to the challenges for current meshing techniques, e.g., blocking technique (Mao et al. 2013b) to capture anatomical details, while is also a reasonable trade-off for computational efficiency. However, lacking these anatomical details hinders a model’s capacity for studying certain localized injuries such as at sulci, gyri and surrounding ventricles (further discussion found below). Nevertheless, these models with high computational efficiency have played important roles in improving our understanding of TBIs; some have found wide applications for improved vehicle safety and helmet design. Modeling techniques learned from the these head models also pave the way for future models with higher anatomical accuracy.

To address anatomical accuracy, voxel-based approach has been used to generate head models including detailed sulci, gyri, and ventricles (Ho and Kleiven 2009; Chen and Ostoja-Starzewski 2010; Miller et al. 2016; Ghajari et al. 2017). Voxel-based approach by converting voxels to hexahedral elements directly or with various smoothing algorithms is efficient and has been used widely for FE analysis of bone structures. However, a known concern is the less accurate peak strain/stress predicted from such models, especially on the surfaces due to jaggedness (Viceconti et al. 1998; Samani et al. 2001). Although the jaggedness could be reduced with various smoothing algorithms, e.g., (Camacho et al. 1997; Boyd and Müller 2006), with a larger smoothing factor, which, however, is at the expense of decreased element quality. Similarly, brain strains predicted from voxel-based head models may also have accuracy issues at jagged surfaces of outer CSF-brain and ventricle-brain interfaces, but the accuracy level is unknown and yet to be studied. Nevertheless, careful choice of result analysis, e.g., evaluating overall regional brain strains or strain distributions, allows such models to provide valuable insights attributed to its anatomical accuracy, such as high strains at sulci depth (Ho and Kleiven 2009; Ghajari et al. 2017), in line with an earlier experimental study (Lauret et al. 2009). Integrating neuroimaging with model-predicted brain strains has provided a possible association between mechanical response and chronic traumatic encephalopathy (CTE) (Ghajari et al. 2017). However, when brain strains at the jagged interfaces are of primary interest, models with conforming meshes capturing sulci, gyri, and brain ventricles are preferred. The jagged interfaces also hinder a reliable implementation of sliding or fluid-structure interaction (FSI). Lastly, falx and tentorium also need to be manually generated (Ho and Kleiven 2009; Miller et al. 2016), affecting its subject-specific efficiency; some smooth-voxel model chose not to include falx/tentorium (Chen and Ostoja-Starzewski 2010).

Another technique for efficient generation of subject-specific models is by mesh morphing (also called warping). The concept has been used extensively in many biomechanics fields on different organs (Couteau et al. 2000; Castellano-Smith et al. 2001; Fernandez et al. 2004; Sigal et al. 2008; Bucki et al. 2010; Bijar et al. 2016; Park et al. 2017), full-body models (Davis et al. 2016; Beillas and Berthet 2017; Liu et al. 2020), as well as smooth brain models (Hu et al. 2007; Ji et al. 2011; Ji et al. 2015; Wu et al. 2019), showing promising results. A typical procedure includes image registration (rigid/affine and/or followed by nonlinear registration algorithms), from which displacement field representing the geometrical difference between the subject and baseline mesh is obtained. Next, the displacement field is applied to morph the baseline mesh, resulting in a personalized mesh with updated nodal coordinates while remaining element connections. In general, the computed displacement field should comply with continuum mechanics conditions on motion, requiring diffeomorphic, non-folding, and one-to-one correspondence to avoid excessive element distortions (Bucki et al. 2010). Otherwise, without such reasonable element quality, not only an FE analysis is prevented from being carried out, also numerical accuracy is influenced. Morphing an anatomically detailed head model that has refined mesh sizes poses a higher requirement on smoothness (associated with Jacobian) of the computed displacement field, meanwhile provides an opportunity attributing to FE model’s direct correspondence with neuroimaging and allows utilizing the advanced registration algorithms developed within the neuroimaging field. Therefore, although mesh morphing is efficient, one major challenge for using it to generate detailed subject-specific FE head model is how to design an image registration pipeline that leads to a displacement field representing well the inter-subject difference of local brain structures, meanwhile not cause excessive element distortions.

Note that anatomical accuracy for a head model is only one among the several factors influencing its biofidelity; other factors include material properties, representation of interactions between various intracranial components (e.g., brain–skull interface). Nevertheless, an anatomically accurate model provides a prerequisite for capturing local strains in areas of interest. Further, an anatomically accurate model due to a representation of the anatomical information from medical images, a direct subject-specific analysis of brain components can be done with higher accuracy compared with coarse mesh models that are made as indirect anatomical representations with reasonable volumetry but not the same mm-accurate anatomical details. Generation of anatomically accurate and subject-specific head models with conforming hexahedral meshes remains a significant challenge based on above literature review. Though conforming tetrahedral meshes are relatively easier to generate, it’s not preferred in head models intended for studying TBIs due to known unfavorable characteristics, such as overstiffening and volumetric locking especially with incompressible material using first order tetrahedral element; though second order could alleviate but may lead to a larger computational cost than hexahedral meshes (Samani et al. 2001).

Despite promising progress, detailed TBI mechanisms remain largely unknown, reflected by not able to predict clinical symptoms, and individual-specific injury tolerances may explain to some extent (Rowson et al. 2018). Subject-specific models for more detailed mechanics of brain injuries are needed. Yet, how brain morphology and WM fiber tract morphology differences among individuals may influence brain injuries remain unclear, although a previous study investigated influences of brain sizes by global scaling (Kleiven & von Holst 2002) and inter-subject WM fiber tract influence by inserting subject’s WM to the same generic head model (Giordano et al. 2017). Head size and shape vary significantly among individuals, as well as WM fiber tracts (Giordano et al. 2017), how these together may influence the brain strain responses are yet to be studied.

This study attempts to address the two challenges: (1) To develop an anatomically detailed head model with conforming hexahedral meshes; (2) Takes inspiration from pioneer works on mesh morphing and develop a new image registration pipeline for morphing a detailed head model. To address the first, the meshing approach used in our previous studies (Li et al. 2017; Li et al. 2019; Zhou et al. 2019b; Zhou et al. 2020) is used. Especially the approach has been shown to generate a detailed elderly head model with a smooth interface between the brain and CSF, permitting a successful implementation of FSI at ventricle-brain interface for studying periventricular injury (Zhou et al. 2020). Efforts toward the above two directions lead to the development of the Detailed and Personalizable Head Model with Axons for Injury Prediction (defined as the ADAPT head model), and equally important a hierarchical image registration pipeline for detailed subject-specific head model generation by morphing. The ADAPT head model is an anatomically detailed head model, including sulci, gyri, connecting ventricular system with conforming mesh, and embedded with WM fiber tracts. The hierarchical pipeline integrating Demons and Dramms deformable registration leads to personalized (i.e., subject-specific) models with satisfactory element quality without further mesh repairing. The uniqueness of the ADAPT head model is the equipped pipeline that allows fast generation of detailed subject-specific models with large variations in head size/shape as well as local brain regions and lateral ventricles with competitive personalization accuracy. The research community may find the hierarchical image registration pipeline useful to morph other head models as well, such as smoothed-voxel head models.

This study is organized as below: Firstly, the development and validation of the ADAPT head model is presented. Secondly, the hierarchical image registration pipeline for personalization is described and its capacity is exemplified by generating six subject-specific head models with largely varying intracranial volumes (ICVs) and brain shapes. Personalization accuracy is quantified by DICE similarity coefficients. Lastly, we use the six subject-specific head models to study the influences of brain size/shape on brain strain response under the same concussive impact. We hypothesize that a large variation in brain strain and location may exist among subjects and could be revealed by anatomically detailed subject-specific models.

## 2 Method

### 2.1 Head model development

The geometry of the ADAPT head model is based on reconstructions of the ICBM152 template generated from 152 healthy subjects (18–43.5 years) (Fonov et al. 2011; Fonov et al. 2009), including T1W, T2W images, and probability maps. T1W and T2W images are segmented using an Expectation-Maximization (EM) algorithm together with the spatial information provided by the probability maps using the software *Slicer 3D* (3D Slicer Version 3.6 2010; Fedorov et al. 2012). Three-dimensional (3D) triangular surface meshes are then generated based on the segmented images and serve as input to the software *Hexotic* to generate all hexahedral elements using an Octree-algorithm (Maréchal 2009). The head model includes the brain, skull (compact and diploe porous bone), meninges (pia, dura, falx, and tentorium), CSF, and superior sagittal sinus (SSS) (**Fig. 1**). The brain is divided into primary structures of cerebral gray matter (GM) (i.e., cerebral cortex), cerebral white matter (WM), corpus callosum (CC), brain stem (BS), cerebellum GM and WM, thalamus, and hippocampus. CSF are divided into outer CSF and ventricular system including lateral ventricles, 3^rd^ and 4^th^ ventricles connected by cerebral aqueduct. Continuous mesh is used throughout the model, with all meshes node-connected from the brain, pia, CSF and dura to the inner skull, including all interfaces between the outer CSF and the brain near sulci and gyri. Note as the same material property is used for the entire brain, dividing it into subcomponents is mainly for post-processing purposes. Further, a maximum level of recursive partitioning on the initial octree cube is set to eight in the software *Hexotic* during meshing to allow capturing the complex structures of sulci, gyri, and ventricular system, resulting in smallest element size of about 0.5 mm at these areas, and transits to larger-sized elements to 1 mm and largest about 2.5 mm at inner brain areas as shown in **Fig. 2**b. A smooth element size transition is ensured by the *balancing rule* implemented in *Hexotic*, with details found in Maréchal (2009). The total number of elements in the head model is 4.4 million hexahedral and 0.54 million quad elements. The minimum Jacobian in the brain is 0.45. All simulations are conducted with LS-Dyna 971 R11 using an explicit dynamic solving method. A typical impact loading with a duration of 100 ms takes about 22 hours using massively parallel processing version with 256 CPUs.

**Fig. 1.**
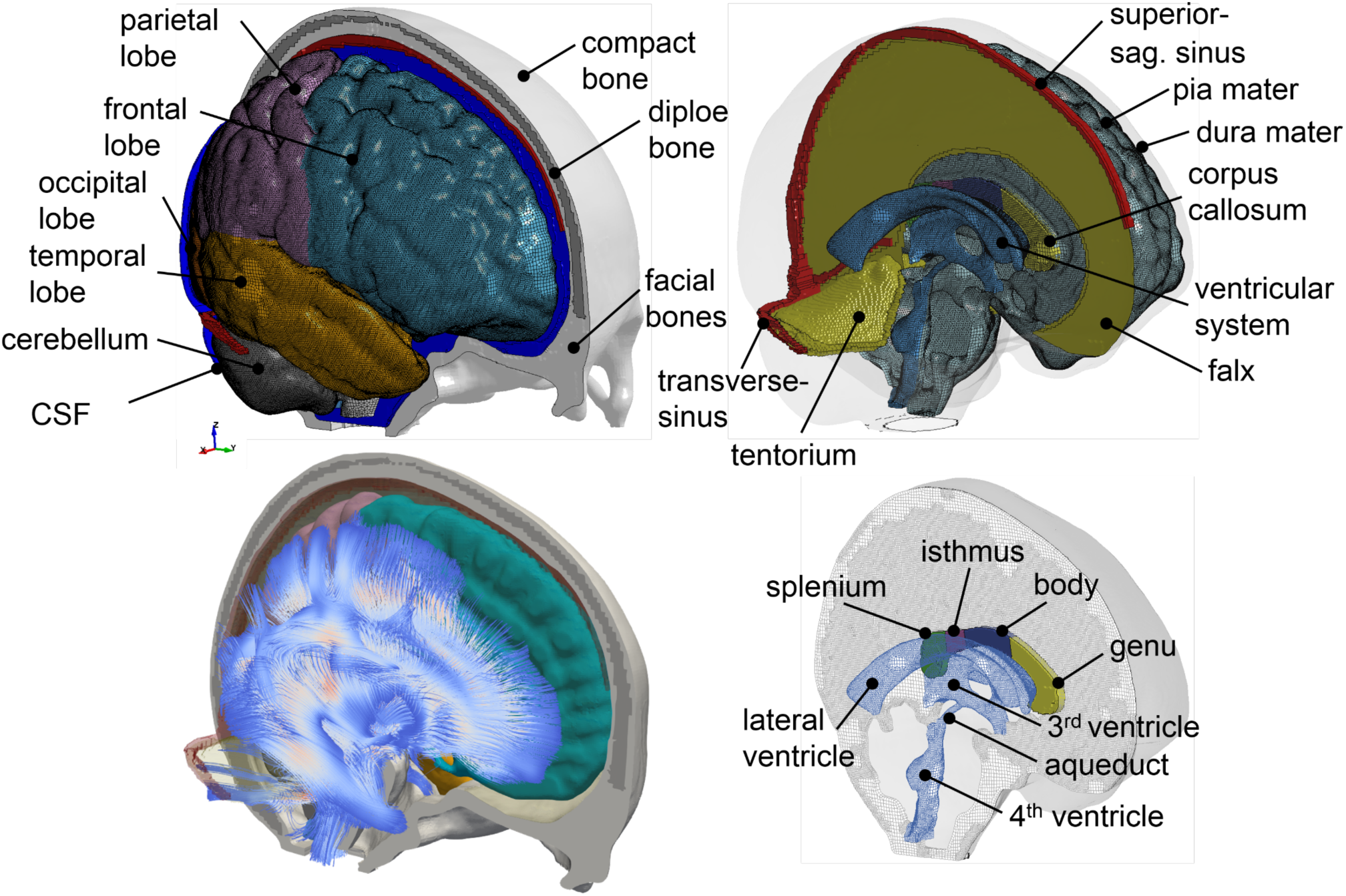
The ADAPT head model with major components illustrated (upper), embedded with WM fiber tracts (lower left), and with connecting ventricular system to the outer CSF (lower right).

**Fig. 2.**
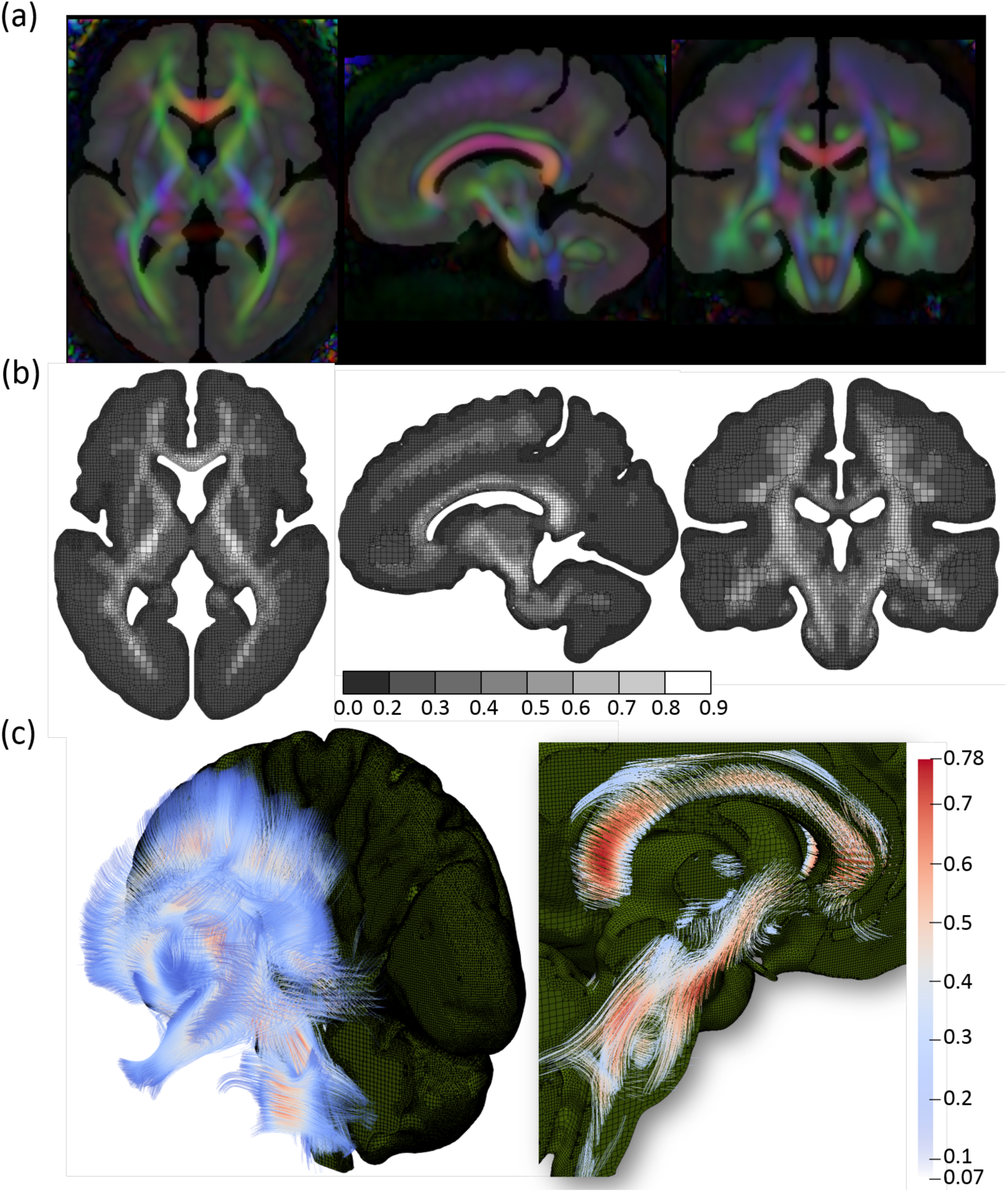
Mapping DTI to the FE head model. (a) Brain mask image corresponding to the FE model overlay with FA calculated from DTI, showing DTI information is directly mappable to the FE model without geometrical adaption. Color-coded FA reflects the WM orientation (red [right–left)], green [anterior-posterior], and blue [superior– inferior)]). (b) Brain FE element with FA mapped. (c) View of the brain embedded with axonal fiber tracts with an enlarged image showing the axonal fibers at CC and BS with FA color scale indicated.

The brain is modeled as hyper-viscoelastic material to account for large deformations of the tissue, with additional linear viscoelastic terms to account for the rate dependence. Material properties presented by Kleiven (2007) is used, which was based on careful analysis of experimental data. Pia, dura/falx/tentorium are modeled with nonlinear hyperelastic material using simplified rubber/foam based on the average stress-strain experimental data (Aimedieu and Grebe 2004; Van Noort et al. 1981). Material constants used in all parts of the head model are summarized in **Table 1**.

**Table 1.**
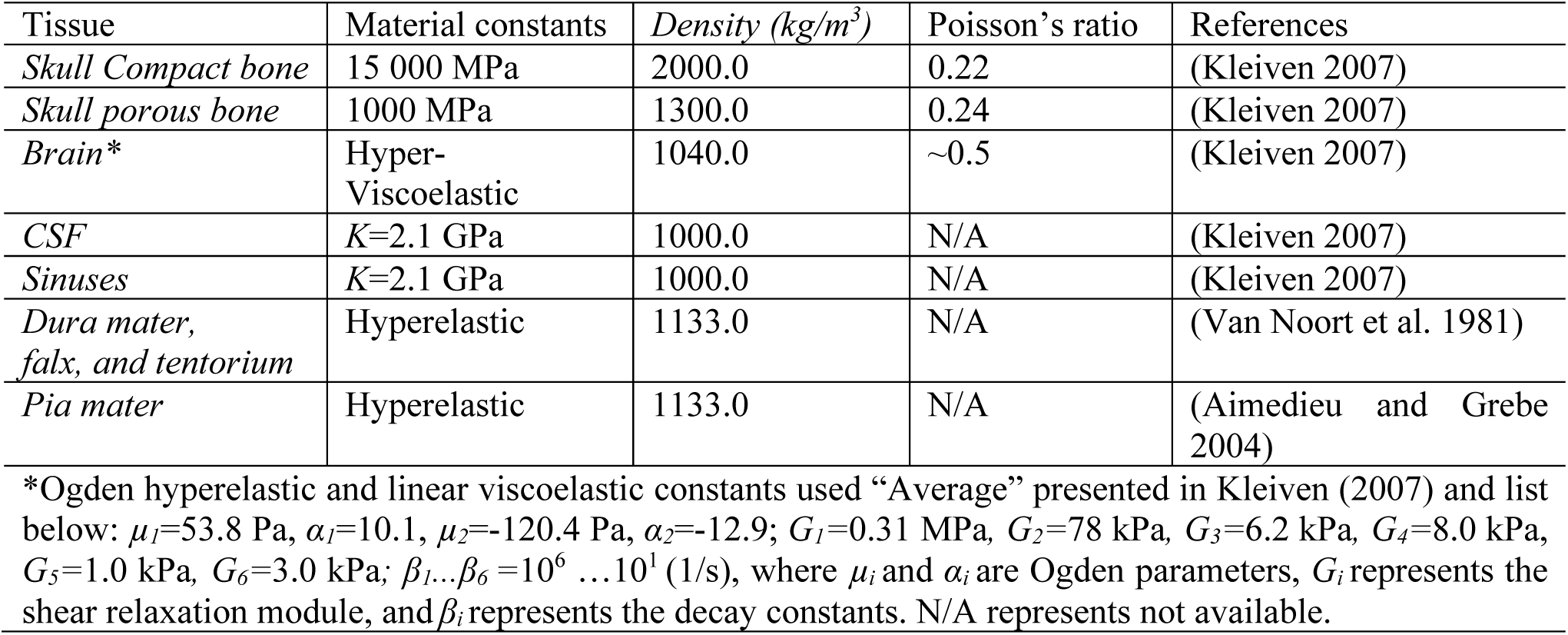
Material properties used in the head model.

### 2.2 Map DTI into the head model for axonal strain calculation

The ADAPT head model is embedded with WM fiber tracts extracted from the ICBM DTI-81 atlas (Mori et al. 2008), which contains white matter information fused with the ICBM152 template space. Eigenvalues and eigenvectors of the already calculated diffusion tensors from the ICBM DTI-81 atlas are calculated at each voxel, based on which Fractional Anisotropy (FA) and WM fiber tracts are obtained. Briefly, FA is calculated as a normalized expression of the eigenvalues and streamline method (Mori et al. 1999; Mori and Zhang 2006) is then used to extract WM fiber tracts associated with the 1^st^ principal eigenvector. A more detailed description of FA and fiber tract extraction can be found in earlier studies (Li et al. 2013; von Holst and Li 2013). The calculated FA values at each voxel with a resolution of 1 mm is shown in **Fig. 2**a.

To calculate axonal strain as defined in Eqn.1, DTI information needs to be mapped to the FE head model. As the geometry of the ADAPT head model is based on the same template as DTI, diffusion tensors from the ICBM DTI-81 atlas are directly mappable to the FE head model without geometrical adaption. Briefly, the DTI voxel closest to the centroid of each FE element is identified based on their spatial coordinates, and the FA and 1^st^ principal eigenvector for this voxel is linked to each FE element. The resultant FA mapped at FE brain resolution is shown in **Fig. 2**b. The final extracted WM tracts in the whole brain contain polylines aligned with the FE head model is shown in **Fig. 2**c (left), from which the CC and BS fiber tracts are enlarged (**Fig. 2**c right). Note the embedded WM fiber tracts are not used in this study, rather diffusion tensors extracted from the subject’s own diffusion-weighted imaging (DWI) are mapped directly to the subject-specific head models (see Sec. 2.5.2). Nevertheless, the paired WM fibers tracts to the baseline head model is useful for future studies when subject’s DTI is not available.

Green-Lagrange strain in the direction of WM tract (abbreviated as axonal strain hereafter) is obtained by projecting calculated strain tensor of each element extracted from LS-Dyna solver along the axonal fiber direction according to the following equation (Giordano et al. 2014):

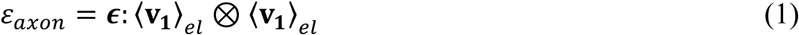

where ***ϵ*** represents the Green-Lagrange strain tensor of each element in a Cartesian vector basis and〈**V**_**1**_〉_*el*_ denotes the axonal fiber direction in the same element obtained as the 1^st^ eigenvector of the diffusion tensor.

### 2.3 Validation performance of the head model and CORA calculation

The performance of the head model is evaluated by comparing experimental data of brain-skull relative motion, brain strain, as well as intracranial pressure close to or at injury level. For all the selected validation experiments mentioned above, the model is scaled to match the anthropometric measurement of the cadaveric heads. To further evaluate whether the model could predict brain response under noninjuries level in living subjects, brain-skull relative motion and brain strain are compared with the experimental displacements and strains measured in a human volunteer using tagged MRI during mild frontal impact presented in Feng et al. (2010). All details of the validation setup are presented in **Supplementary Material**, with a brief description provided below.

For brain-skull relative motion validation, neutral density target (NDT) displacement curves from seven representative cases from Hardy et al. (2007) are selected, including one sagittal (C288-T3), one horizontal (C380-T2), and five coronal impacts (C380-T1, C380-T3, C380-T4, C380-T6, and C393-T3). The recalculated cluster brain strains of these seven selected cases presented in Zhou et al. (2019b) are used as experimental strain data to evaluate brain strain performance of the model. Cluster brain strains from the above seven cases are chosen, with further motivation provided in Discussion. Intracranial pressure response of the head model is compared with recordings from experiment No. 37 conducted by Nahum et al. (1977).

CORA (CORrelation and Analysis, version 3.6.1) scores are calculated to assess the level of correlation between a pair of time history curves using a sub-method included in CORA, i.e., the cross-correlation method. CORA score reported in this study is calculated as (V+G+P)/3 in terms of shape (V), size (G), and phase (P), meaning equal weights for the three parts. CORA scores range from 0 to 1 with 1 indicating a perfect match. Note another sub-method included in CORA, i.e., the corridor method is excluded, and recommended settings from (Giordano and Kleiven 2016) are adopted in this study with details provided therein.

### 2.4 Hierarchical image registration pipeline for mesh morphing

The personalization approach for subject-specific head model generation is based on Demons and Dramms deformable registrations (**Fig. 3**a-i). First, the diffeomorphic Demons registration (Vercauteren et al. 2009) implemented in the open-source software *Slicer 3D* is performed between the segmented cranial masks of the baseline (corresponding to the baseline FE mesh) and the subject after being rigidly aligned using a 6 degree-of-freedom rigid registration available in *Slicer* 3D. Further details for the Demons registration steps can be found in a previous study (von Holst and Li 2013). Afterward, Dramms registration algorithm (Ou et al. 2011) implemented as open-source code by the authors (Dramms version 1.5.1, 2018) is performed on the skull stripped T1W images inherited from the Demons step. The resultant displacement field from the two-step registrations is then applied to morph the baseline mesh of the ADAPT model, obtaining a subject-specific model. The subject’s own DTI is then mapped to the personalize model using the same procedure described in Sec. 2.2 resulting in a subject-specific model incorporated with the subject’s own WM fiber tracts (**Fig. 3**j).

**Fig. 3.**
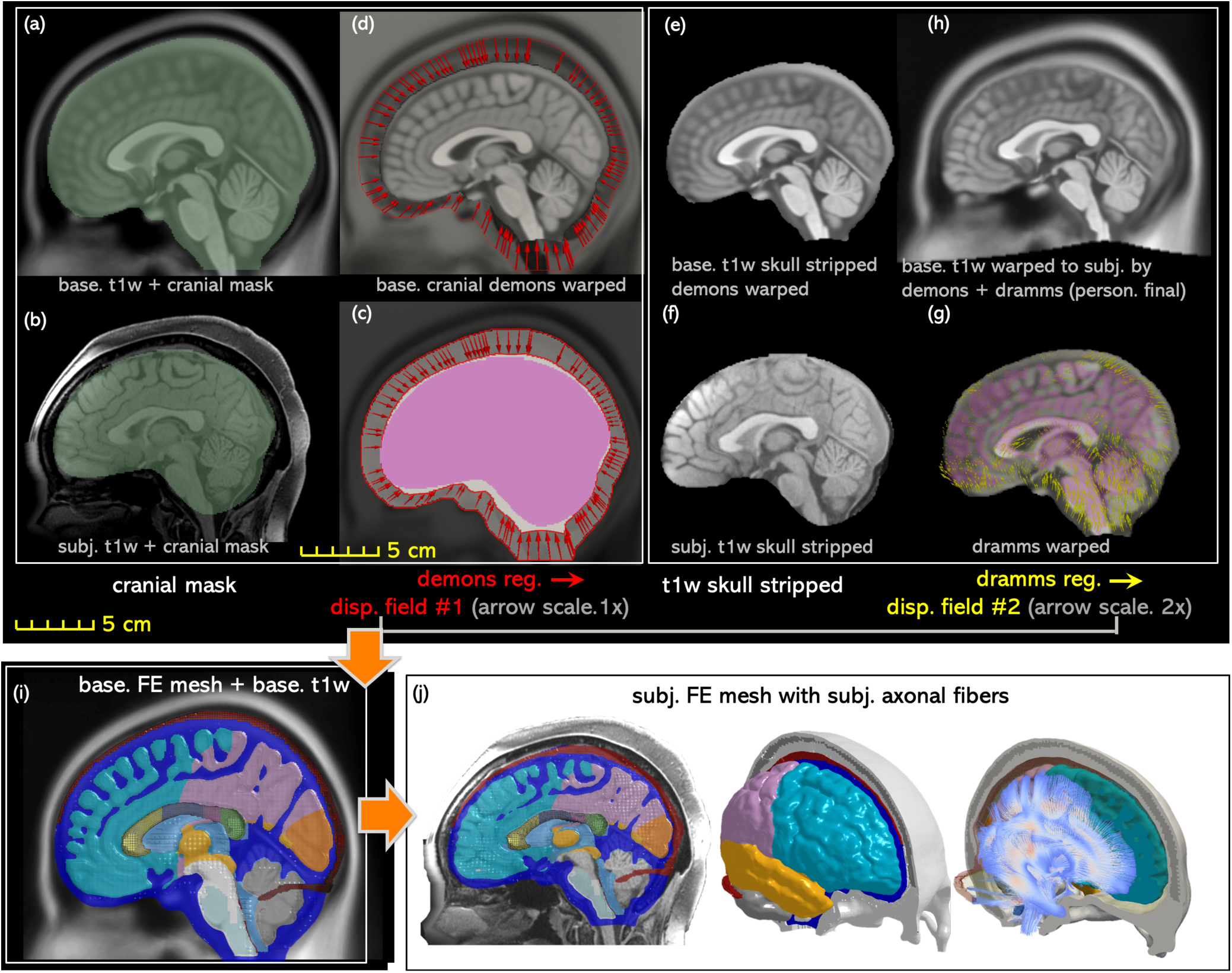
The workflow of the proposed hierarchical image registration pipeline for subject-specific head model generation by morphing demonstrated with the results from the smallest female. Baseline T1W image (i.e., the T1W image corresponding to the baseline FE mesh) and the subject’s T1W are segmented to obtain the cranial mask (a, b), which are used as input for Demons registration from which displacement field #1 is obtained as indicated by the arrows (c). Displacement field #1 is further applied to the baseline T1W image (d), which is then skull stripped (e) and afterward together with the subject’s skull stripped T1W (f) as input to Dramms registration (g). The obtained displacement field #2 from Dramms registration is applied to the baseline T1W and obtain the personalized T1W (i.e., the T1W image corresponding to the subject-specific mesh), which is compared with subject’s T1W to evaluate personalization accuracy. Finally, the two displacement fields add up to morph the baseline mesh (i), obtaining the subject-specific head model, including both the mesh and WM fiber tracts (j).

### 2.5 Subject-specific models with axonal fibers

The capacity of the personalization approach is demonstrated by generating six models with largely varying ICV. First, six subjects are identified by analyzing the ICVs from the WU-Minn Human Connectome Project (WUM HCP) database (Van Essen et al., 2013). For all the six subjects, both T1W and DWI images are all openly accessible. T1W image is used for subject-specific mesh generation, and DWIs are mapped to the personalized model, with details presented below.

#### 2.5.1 Six subjects identified from HCP

Out of the 1200 subjects from the WUM HCP database, 3T structural scans of T1W are available for 1113 subjects, and the data of ICV are already processed using the software FreeSurfer (Glasser et al., 2013). As listed in **Table 2**, six subjects covering a wide range of ICV are selected, including the smallest head (turns out to be a female), the largest head (turns out to be a male), and four heads with ICVs in between.

**Table 2.** Subjects selected from the HCP database representing heads of largely varying ICV.

| Description | Subject ID | Age group | ICV (dl) |
| --- | --- | --- | --- |
| Smallest (female) | 118124 | '31-35' | 831.21 |
| 5<sup>th</sup> perc. female | 568963 | '31-35' | 1186.1 |
| 50<sup>th</sup> perc. female | 771354 | '26-30' | 1479.6 |
| 5<sup>th</sup> perc. male | 185038 | '31-35' | 1474.0 |
| 50<sup>th</sup> perc. male | 172635 | '31-35' | 1697.3 |
| Largest (male) | 223929 | '31-35' | 2143.2 |

Note: The 5<sup>th</sup> percentile male and the 50<sup>th</sup> percentile female have nearly the same ICV.

#### 2.5.2 WM fiber tracts extracted from DWIs

The DWIs of the six subjects with a resolution of 1.25 mm are processed to extract WM fibers. The downloaded DWI dataset has already been preprocessed, including corrections for gradient non-linearity, motion-correction, and eddy-currents (Glasser et al., 2013), which are further processed to extract diffusion tensors and WM fiber direction (i.e., 1^st^ eigenvector) in each voxel using FSL v6.0.2 DTIFit with a weighted linear least squares option. The WM fiber directions are then mapped directly to the subject-specific mesh using the approach described in Sec.2.2, based on which axonal strains are calculated.

### 2.6 Personalization accuracy evaluation

The baseline T1W image is morphed to a personalized T1W image for each subject (see **Appendix 1**) using the procedure presented in Sec. 2.4. DICE is then calculated to quantify personalization accuracy, i.e., how well the personalized T1W image (corresponding to the subject-specific mesh) reflects the subject’s T1W as the ground truth. DICE is a single metric commonly used in neuroimaging field (Bennett and Miller 2010; Zou et al. 2004) to measure the spatial overlap between two segmentations (A and B), and is defined as DICE(A,B) = 2(A∩B)/(A+B) where ∩ is the intersection. DICE value of 0 implies no overlap at all between both, whereas a DICE coefficient of 1 indicates perfect overlap.

To calculate DICE, automated segmentation is performed using the software FreeSurfer (version 7.1.0) with the default brain segmentation pipeline (*recon-all*) for both the personalized and subjects’ T1W images without additional manual editing. For CC, one midsagittal plan is segmented by thresholding and used for DICE calculation instead of using FreeSurfer segmented CC due to inaccuracy using the default pipeline. Segmented labels are combined to the following components for DICE evaluation: brain, local brain regions of cerebral GM & WM, CC, BS, hippocampus, thalamus, and cerebellum, as exemplified with two subjects (**Fig. 4**). Note DICE value for cranial mask is calculated based on the segmented image in *Slicer 3D* as shown in Fig. 3a &b, instead of using FreeSurfer segmented due to insufficient quality of the skull-stripped cranial using *recon-all*. Also note that FreeSurfer segmented labels are only used during DICE calculation, and the quality of the automatic segmentation has no influence on the subject-specific mesh development process.

**Fig. 4.**
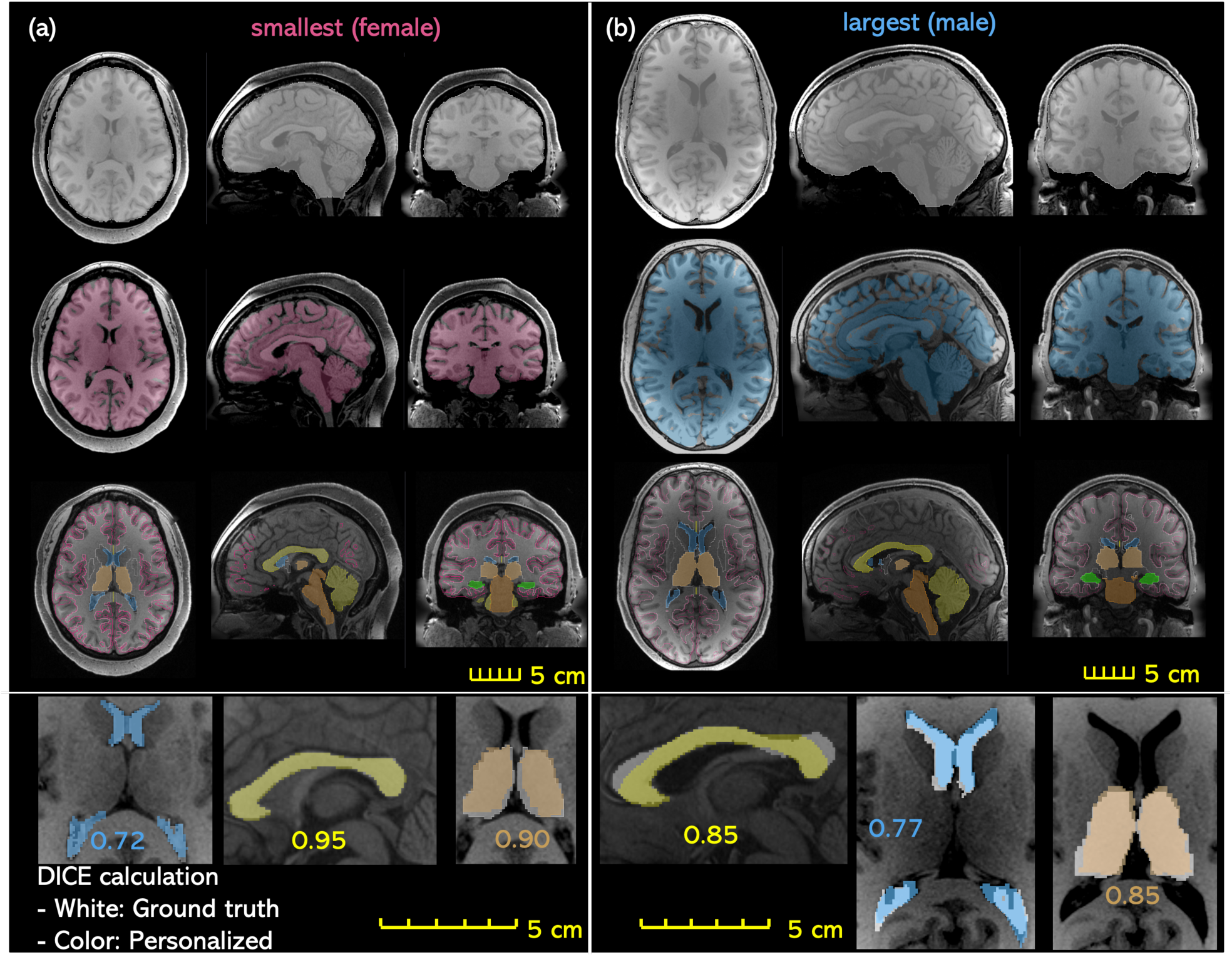
Evaluation of DICE exemplified with the smallest (a) and the largest head (b), including global structures of the cranial mask (row 1), the brain (row 2), local brain regions of cerebrum GM, WM, cerebellum, hippocampus, thalamus, CC, BS, as well as lateral ventricles. T1W image of the subject is overplayed with the segmented regions from the personalized T1W image (row 3). Enlarged figures show the segmented regions from the personalized T1W image (shown in color) is overlayed with the segmented regions from subject’s T1W as ground truth (gray), based on which DICE is calculated as exemplified for lateral ventricles, corpus callosum, and thalamus with DICE values shown (row 4).

### 2.7 Loading conditions and brain strain evaluation

The six subject-specific head models are loaded with the same impact kinetics measured in a collegiate American football player resulting in loss of consciousness reported earlier (Hernandez et al. 2015). Translational accelerations and rotational accelerations (**Fig. 5**) are imposed on the center of gravity (C.G) of the head models.

**Fig. 5.**
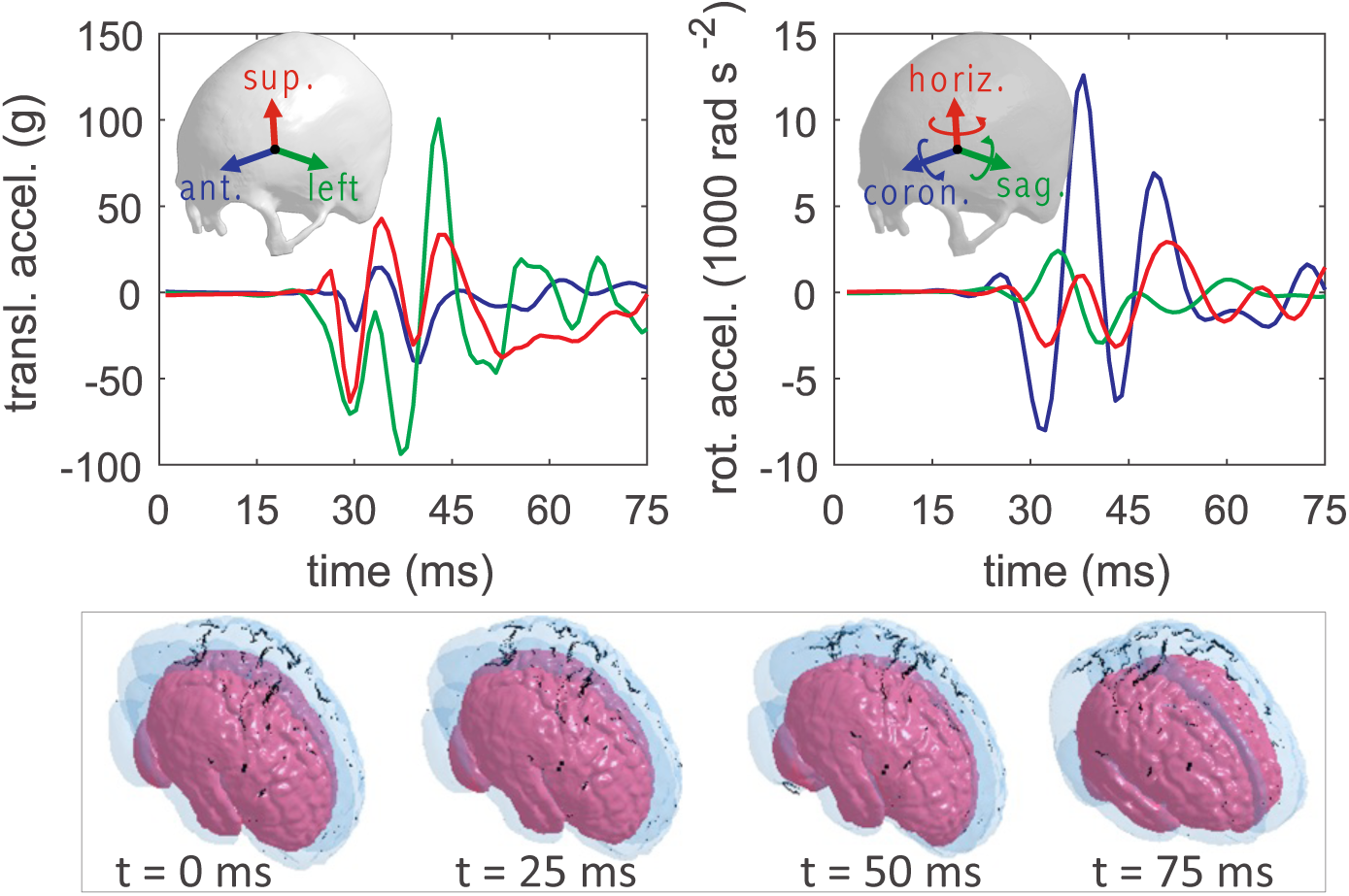
Translational (upper left) and rotational accelerations (upper right) loaded to the six subject-specific head models. All the six models are subjected to the same loading, and the head positions during the impact in two subjects are illustrated (lower row), with the smallest female in pink color and the largest male in transparent blue (dark stripes relating to rendering only).

Maximum of the 1^st^ principal Green-Lagrange strain (MPS), and maximum axonal strain (MAS) (i.e., strains along the WM fiber direction as defined in Eqn.1) during the entire impact, as well as the locations of both metrics, are analyzed and compared between the six subjects.

## 3 Results

### 3.1 Validation performance of the ADAPT head model

#### 3.1.1 Brain-skull relative motion

The CORA scores for the ADAPT head model on brain motion are presented in **Table 3** in comparison with head models previously developed at the same research group, including the original KTH head model (Kleiven 2007) and its updated version with FSI for brain-skull interface (Zhou et al. 2019a) interface (referred to as KTH-FSI model), as well as KTH detailed head model (Zhou et al. 2019b). Note that, to make the validation performance of different models amenable a direct comparison, the CORA scores for previous models reported here are either newly calculated or recalculated using exactly the same approach and CORA settings described in this study. The CORA scores for the ADAPT head model are higher than the original KTH head model for all the seven cases, resulting a higher mean CORA score being 0.617 versus 0.493, respectively. While CORA scores are comparable with the KTH-FSI model for the two cases evaluated (C288-T3, C380-T4). The mean CORA score for the ADAPT is slightly lower than the KTH detailed head model (mean CORA score 0.655). To further compare predictions between the ADAPT and the original KTH head model, NDT curves of brain-skull relative motion for three representative cases (sagittal C288-T3, horizontal C380-T2, coronal impact C380-T1) comparing with the experimental data (Hardy et al. 2007) are presented in **Appendix 2**. Curves for all the remaining NDTs predicted from the ADAPT head model in comparation with experimental data are presented in **Supplementary Material.** Using the brain-motion based CORA scores, the ADAPT head model would be rated as “fair” according to the same rating scale used earlier (Zhao and Ji 2020).

**Table 3.**
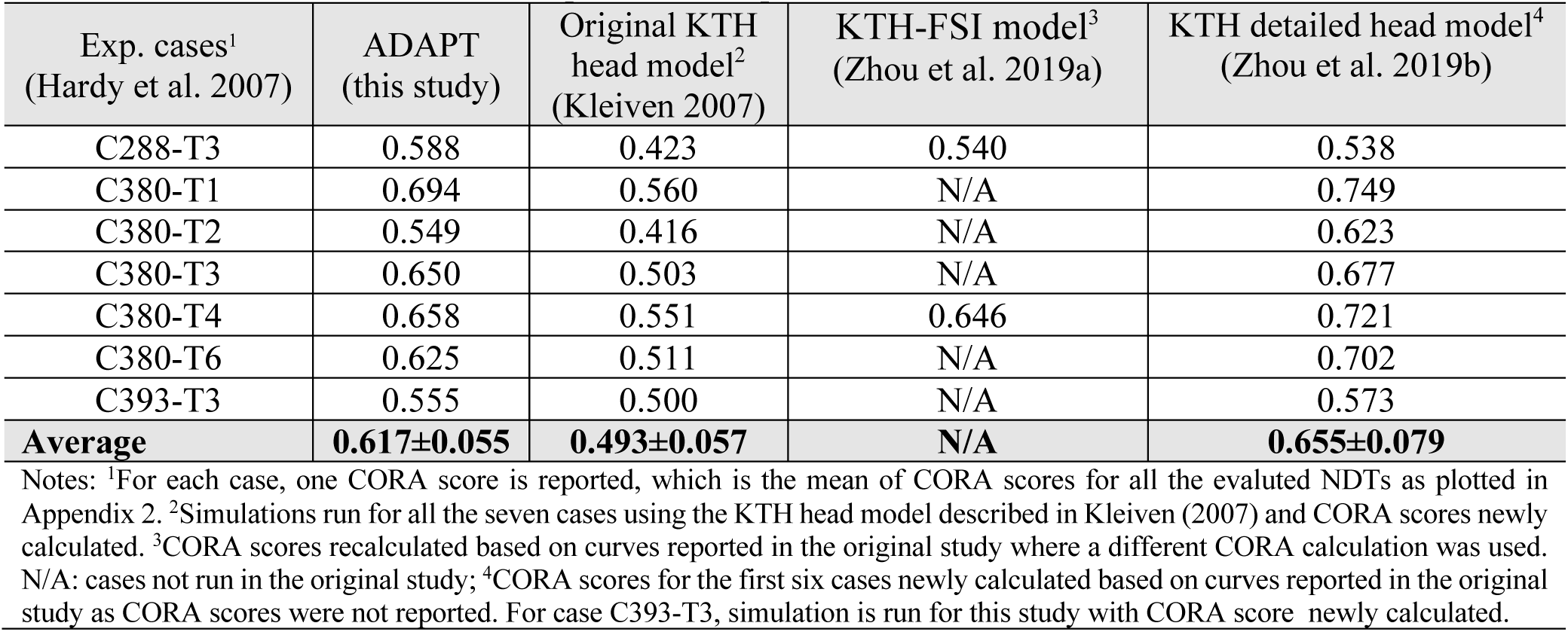
CORA scores for the ADAPT model on brain-skull relative motion in comparation with previous models.

| Exp. cases<sup>1</sup><br>(Hardy et al. 2007) | ADAPT<br>(this study) | Original KTH<br>head model<sup>2</sup><br>(Kleiven 2007) | KTH-FSI model<sup>3</sup><br>(Zhou et al. 2019a) | KTH detailed head model<sup>4</sup><br>(Zhou et al. 2019b) |
| --- | --- | --- | --- | --- |
| C288-T3 | 0.588 | 0.423 | 0.540 | 0.538 |
| C380-T1 | 0.694 | 0.560 | N/A | 0.749 |
| C380-T2 | 0.549 | 0.416 | N/A | 0.623 |
| C380-T3 | 0.650 | 0.503 | N/A | 0.677 |
| C380-T4 | 0.658 | 0.551 | 0.646 | 0.721 |
| C380-T6 | 0.625 | 0.511 | N/A | 0.702 |
| C393-T3 | 0.555 | 0.500 | N/A | 0.573 |
| **Average** | **0.617±0.055** | **0.493±0.057** | **N/A** | **0.655±0.079** |

Notes: <sup>1</sup>For each case, one CORA score is reported, which is the mean of CORA scores for all the evaluated NDTs as plotted in Appendix 2. <sup>2</sup>Simulations run for all the seven cases using the KTH head model described in Kleiven (2007) and CORA scores newly calculated. <sup>3</sup>CORA scores recalculated based on curves reported in the original study where a different CORA calculation was used. N/A: cases not run in the original study; <sup>4</sup>CORA scores for the first six cases newly calculated based on curves reported in the original study as CORA scores were not reported. For case C393-T3, simulation is run for this study with CORA score newly calculated.

#### 3.1.2 Brain strain

The mean CORA scores for the ADAPT head model on principal and shear strain for the seven evaluated clusters are 0.763 and 0.776, respectively (**Table 4**). The CORA scores are compared with the KTH detailed head model (the only model so far that has used the same strain data to systematically evaluate the strain predictablity of FE head model according to the authors’ knowledge) (**Appendix 2, Table A3**), showing comparable values.

**Table 4.**
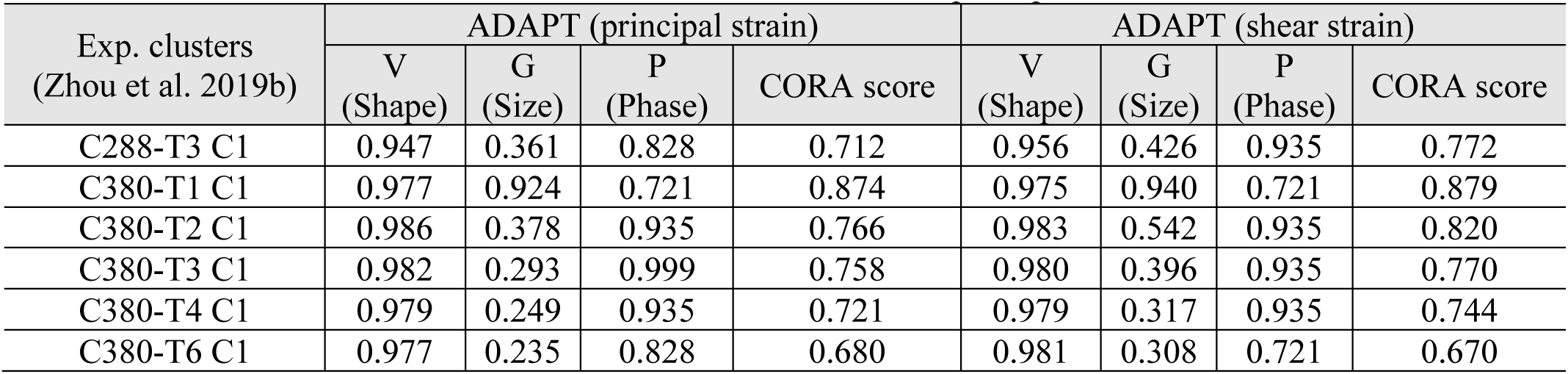
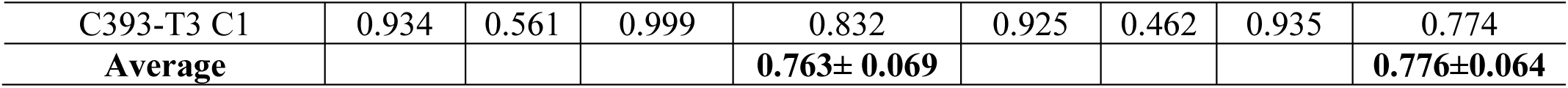
CORA scores for the ADAPT model on principal and shear strain.

| Exp. clusters<br>(Zhou et al. 2019b) | ADAPT (principal strain) |  |  |  | ADAPT (shear strain) |  |  |  |
| --- | --- | --- | --- | --- | --- | --- | --- | --- |
|  | V<br>(Shape) | G<br>(Size) | P<br>(Phase) | CORA score | V<br>(Shape) | G<br>(Size) | P<br>(Phase) | CORA score |
| C288-T3 C1 | 0.947 | 0.361 | 0.828 | 0.712 | 0.956 | 0.426 | 0.935 | 0.772 |
| C380-T1 C1 | 0.977 | 0.924 | 0.721 | 0.874 | 0.975 | 0.940 | 0.721 | 0.879 |
| C380-T2 C1 | 0.986 | 0.378 | 0.935 | 0.766 | 0.983 | 0.542 | 0.935 | 0.820 |
| C380-T3 C1 | 0.982 | 0.293 | 0.999 | 0.758 | 0.980 | 0.396 | 0.935 | 0.770 |
| C380-T4 C1 | 0.979 | 0.249 | 0.935 | 0.721 | 0.979 | 0.317 | 0.935 | 0.744 |
| C380-T6 C1 | 0.977 | 0.235 | 0.828 | 0.680 | 0.981 | 0.308 | 0.721 | 0.670 |
| C393-T3 C1 | 0.934 | 0.561 | 0.999 | 0.832 | 0.925 | 0.462 | 0.935 | 0.774 |
| **Average** |  |  |  | **0.763±0.069** |  |  |  | **0.776±0.064** |

The predicted brain strain curves from the ADAPT model are compared with the experimental brain strain data presented in Zhou et al. (2019b) (**Fig. 6**). Despite a large difference in the peak between the model predicted and experimental data (except for C380-T1 C1 which is closer), the shape and phase show good match, reflected by the high values of V and P; close to 1 in some cases (C380-T3 C1, C393-T3 C1 principal strain) (**Table 4**). Further, the simulated brain strains are consistently lower than the experimental strain in all the seven evaluated clusters except for C380-T1 C1. More clusters are to be studied to see if the same trend holds true for all the 15 cluster brain strains presented in Zhou et al. (2019b). Further discussion on brain strain validation performance and the implications are found in Discussion.

**Fig. 6.**
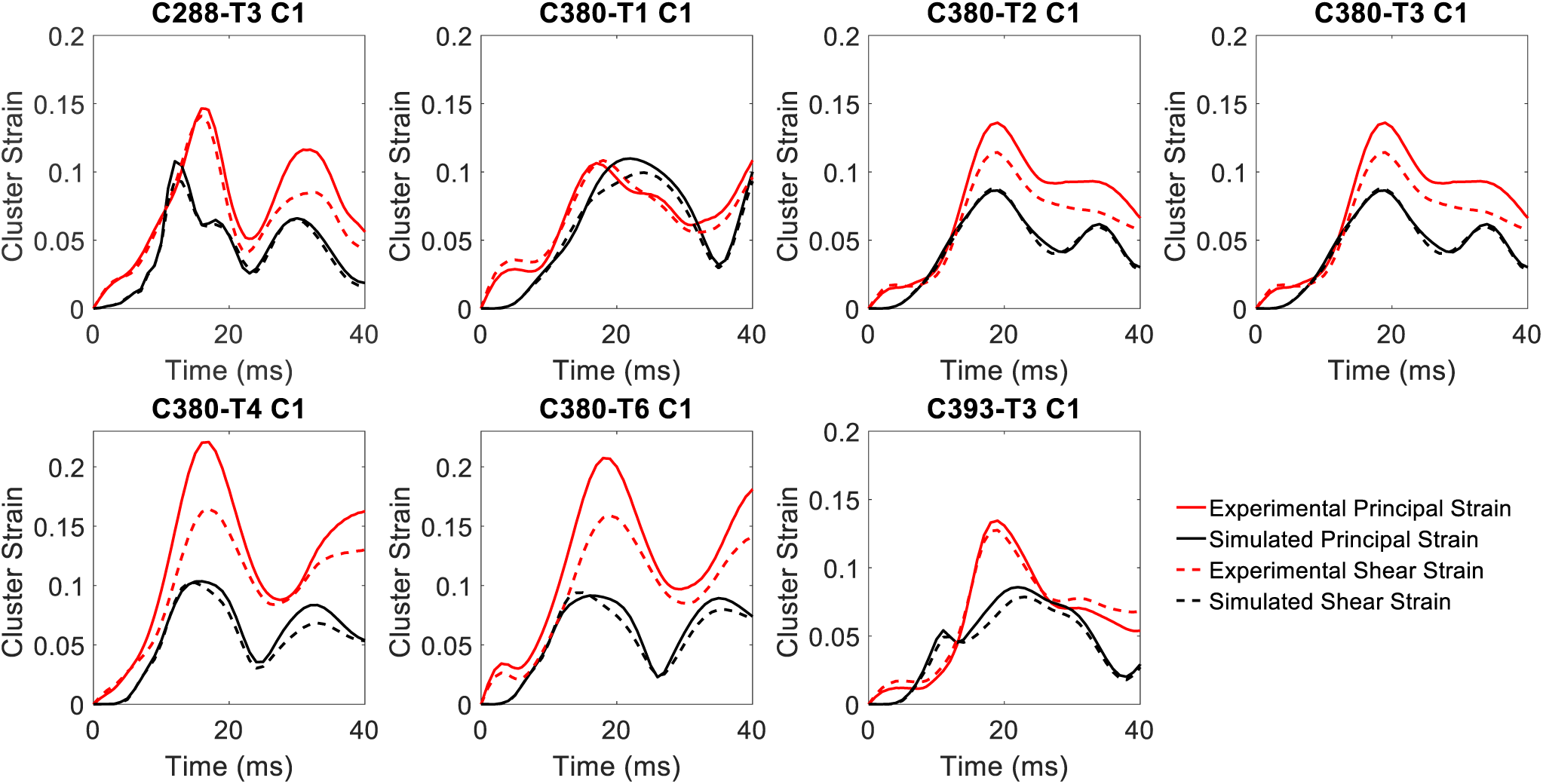
Comparison between the ADAPT model simulated and experimental strain. Experimental strain denotes the recalculated cluster strain presented in Zhou et al. (2019b) based on the original brain-skull relative motion experimental data from Hardy et al. (2007).

The mean CORA scores for brain strain are higher than brain-skull relative motion, attributed to high scores in shape and phase compensating the low values in size (G). Using the brain-strain based CORA scores, the ADAPT head model would be rated as “good” according to the same rating scale used earlier (Zhao and Ji 2020).

#### 3.1.3 Intracranial pressure and in-vivo strain comparison

The CORA scores for the ADAPT head model on intracranial pressure are presented in **Table 5**, with all time-pressure curves presented in **Supplementary Material**. Note the relatively high CORA scores for pressure should not be taken as an overinterpretation of model performance, as multiple studies have shown for an FE model with continuous mesh, brain pressure is uniquely determined by brain mass, brain shape and linear acceleration due to its near incompressibility (Bradshaw and Morfey 2001; Zhao and Ji 2016), as done in cadaveric experiments (Nahum et al. 1977). Nevertheless, a comparation with experimental pressure data may still benefit and serve as additional verification purposes, though it’s important to note brain pressure is less relevant than brain strain for blunt impact simulation.

**Table 5.**
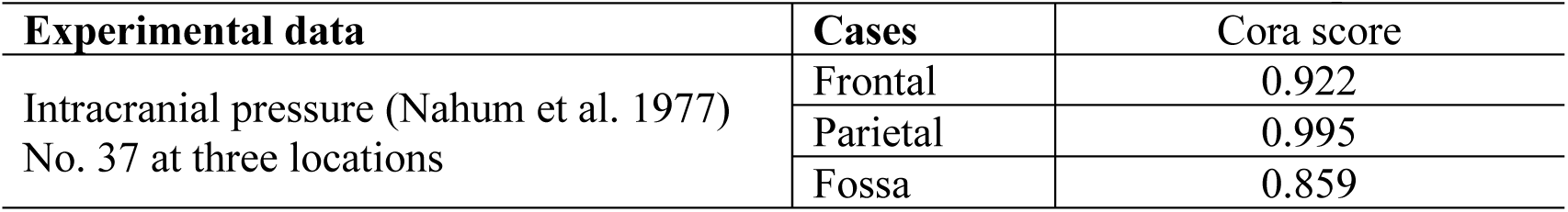
CORA scores of the ADAPT head model on intracranial pressure.

| Experimental data | Cases | Cora score |
| --- | --- | --- |
| Intracranial pressure (Nahum et al. 1977)<br>No. 37 at three locations | Frontal | 0.922 |
|  | Parietal | 0.995 |
|  | Fossa | 0.859 |

The qualitative comparison of brain-skull relative motion and brain strain distribution with in-vivo experimental data (Feng et al. 2010) (presented in **Supplementary** M**aterial**) indeed shows that the current ADAPT model is not capable of predicting the chosen in-vivo measurement. Thus, the comparison shouldn’t be interpreted as the ADAPT model has been validated against such data, rather, to highlight the need and serves as a basis for future investigation.

### 3.2 Personalization accuracy evaluation of DICE for the six subject-specific models

The boxplot of the DICE values of the cranial, the brain, and local brain regions are presented in **Fig. 7** (the DICE values are listed in **Appendix 1)**, in general showing quite good results even for CC with large variations between the baseline and subjects. Especially an average DICE of 0.975, 0.90, and 0.76 are achieved for the cranial mask, the brain, and hippocampus, comparable or even higher than some algorithms used in neuroimaging field (Ou et al. 2014) for capturing inter-subject differences. DICE values for local brain regions, as well as lateral ventricles, are all above 0.6, indicating the internal brain structures of the subject-specific head model reflect the subject to an acceptable level.

**Fig. 7.**
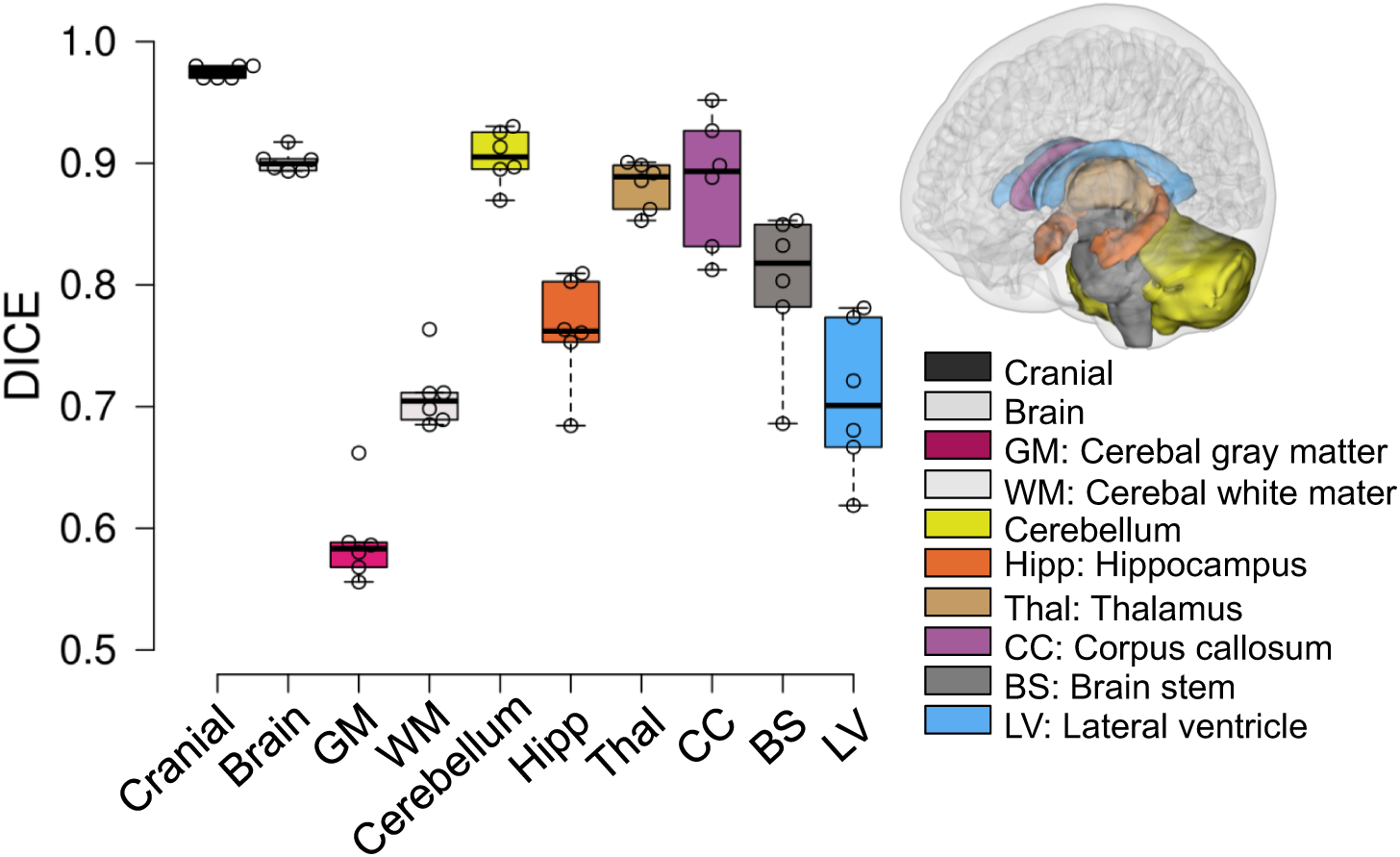
Boxplot of DICE values calculated for the six subjects with regions evaluated shown on the right, including the cranial mask and the brain, local brain regions, and lateral ventricles. The boxplots show the median, minimum, and maximum value.

### 3.3 Subject-specific head model mesh quality

In the six subject-specific head models generated, most brain elements (95.5% ± 1.2% on average for the six subjects) have a Jacobian over 0.5, and almost all elements (99.9%±0.1%) have a Jacobian over 0.45. The minimum Jacobian in the six head models is all above 0.2. In this study, the mesh quality is considered to be satisfactory when at least 95% of the elements have a Jacobian over 0.5. A summary of brain mesh element qualities is listed in **Table A2 (Appendix 1)**.

### 3.4 Magnitude and location of MPS in the cerebral cortex

The time-history curve of MPS in the cerebral cortex is extracted for each subject (**Fig. 8**a). Interestingly, the smallest female shows MPS occurring at 36 ms, slightly different from other subjects except for the largest male occurring at 56 ms (**Fig. 8**a). Note the delay in the peak between both curves is mainly caused by the location difference where maximum strain occurs during the entire impact. The MPS in the cerebral cortex for all subjects is located at the sulci regions, as exemplified with results from two subjects (**Fig. 8**b). Additional animations of the brain strain response during the entire impact for both subjects are provided as **Supplementary Videos**.

**Fig. 8.**
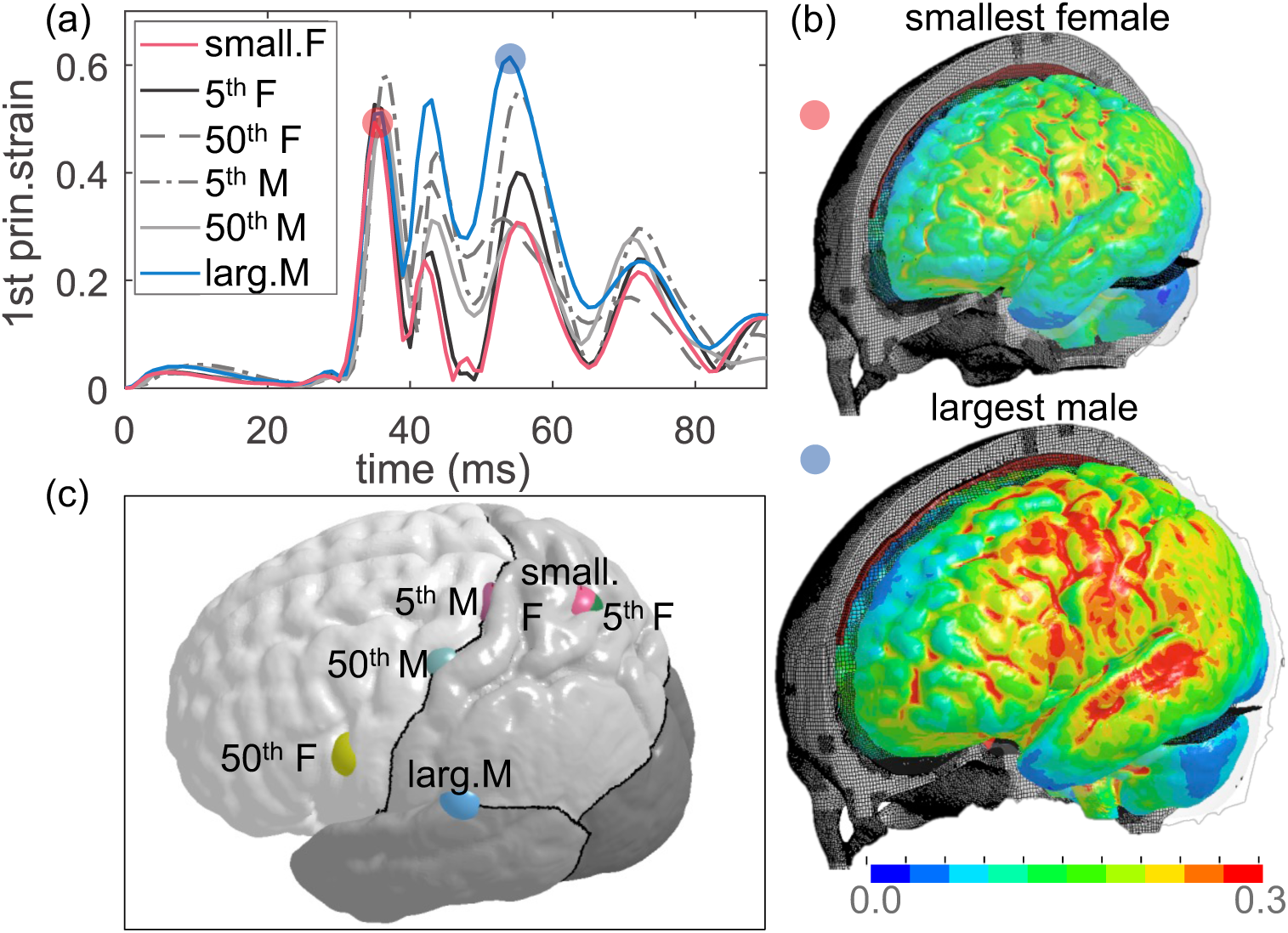
(a) Time-history curves of MPS in the cerebral cortex during the impact of each subject. (b) The strain distribution at the cerebral cortex when MPS occurs exemplified with results from two subjects, captured at 36 ms for the smallest female and 56 ms for the largest male. (c) Locations of MPS shown by a sphere for each subject.

The results show, in general, smaller heads tend to have a lower brain strain under the same impact loading; the lowest MPS (MPS 0.4794) and highest MPS (0.6144) are found in the smallest and largest head respectively. Further, the MPS in other heads with ICV falls within these two cases but does not follow a monotonic trend. The locations of MPS for the six subjects are notably different, shifting between frontal, parietal, and temporal lobes (**Fig. 8**c).

### 3.5 Brain regional response analysis: MPS and MAS

The time-history curves of the 1^st^ G-L principal strain from all elements of cerebral WM, CC, BS, hippocampus, and thalamus are evaluated and exemplified with the results from the smallest and largest heads (**Fig. 9**). The color shaded shape is formed by strain-time history curves from all elements, of which the curve from the element with maximum strain is plotted with black color line for each brain region. Of interest note that MPS typically occurs at a similar time between both subjects in most brain regions (i.e., similar phase in black color line between two models), except for CC and thalamus (**Fig. 9**, 2^nd^ and 5^th^ row), shifting to a later time in the largest head compared with the smallest head. The difference in the shaded shape indicate element-wise different MPS response between the two models, also indicate a different location of MPS. For example, the shade shape for CC and thalamus are notably different between the two subjects, indicating the largest strain likely occur at a different element (location) between the two models at CC and thalamus (similarly as observed for strain at cerebral cortex as shown in **Fig. 8**b).

**Fig. 9.**
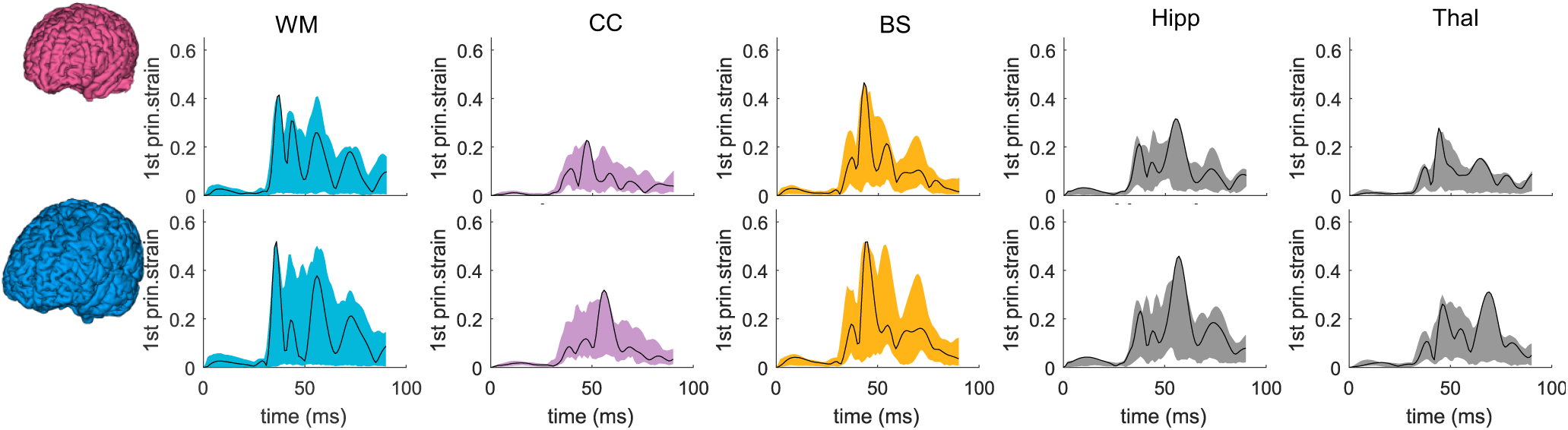
Time-history curves of 1^st^ principal G-L strain for all elements in brain regions of cerebral WM, CC, BS, hippocampus, and thalamus. The color shaded shape is formed by curves from all elements for different brain regions, and the black curve shows the curve in the element with largest strain. Upper row: smallest female. Lower row: largest male.

The results for axonal strain are plotted in **Fig.10** with time-history curves of axonal strain in all elements of cerebral WM, CC, BS plotted. Note axonal strain for thalamus and hippocampus not evaluated (grey matter region with less anisotropy), thus, is not plotted. Similarly, the color shaded shape varies between the two subjects among brain regions, indicating element-wise different MAS response between the two models. The MAS in cerebral WM occurs much later in the smallest female than in the largest male (**Fig. 10**, 1^st^ row), a different trend than observed for MPS with maximum strain occurs at similar time for WM (**Fig. 9**, upper row).

**Fig. 10.**
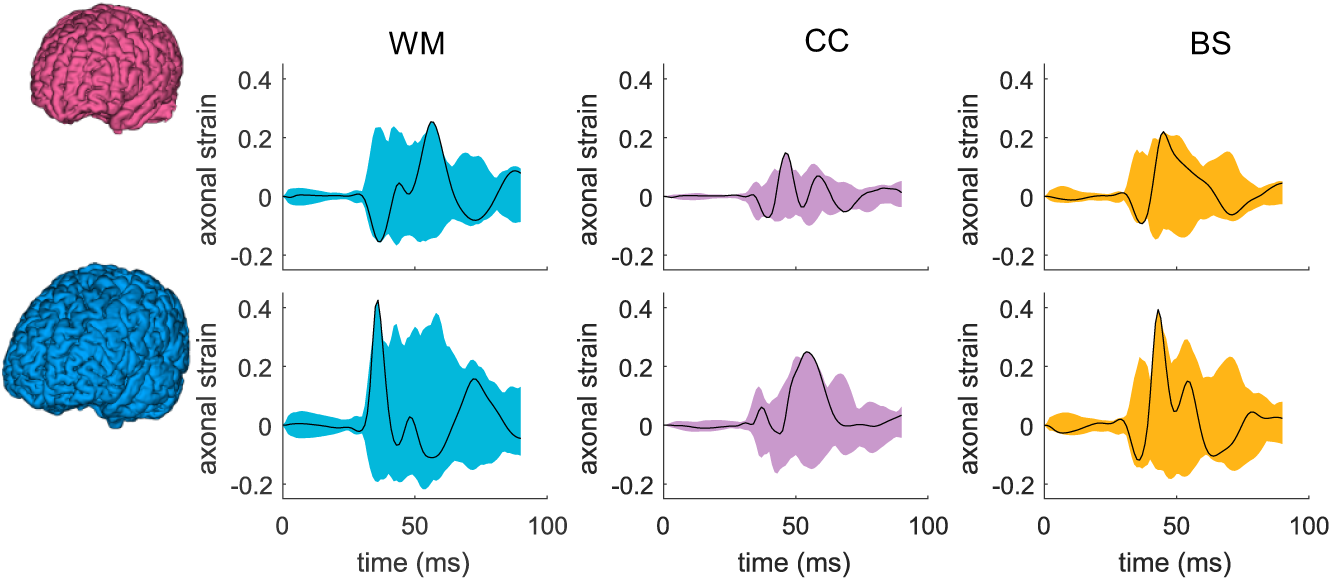
Time-history curves of axonal strain for all elements in brain regions of cerebral WM, CC, and BS. The color shaded shape is formed by curves from all elements for different brain regions, and the black curve shows the curve in the element with largest strain (upper row: smallest female; lower row: largest male).

The time-history curves of all elements for only two subjects are presented above, and for all other subjects, only the maximum values (MPS and MAS) are presented in the boxplot (**Fig. 11**) (exact values are listed in **Appendix 3**). Both the MPS and MAS show significant differences among the six subjects using the one-way ANOVA test (p<0.001 for both MPS and MAS). A similar trend is observed in the cerebral cortex that a smaller brain tends to have a smaller MPS though not following a monotonic trend. In particular, for CC, the largest male has the largest MPS (0.32), compared with the smallest female with the lowest MPS (0.23), differing 40.3%. Of the evaluated brain regions, up to 44.9%, inter-subject differences in MPS are found in the hippocampus. For MAS, up to 86.21% difference in found in CC (the 50^th^ percentile female with the lowest MAS and the largest male with highest MAS). The results show an even larger inter-subject variability in MAS than MPS, which is logical as WM differences are further accounted for when calculating MAS.

**Fig. 11.**
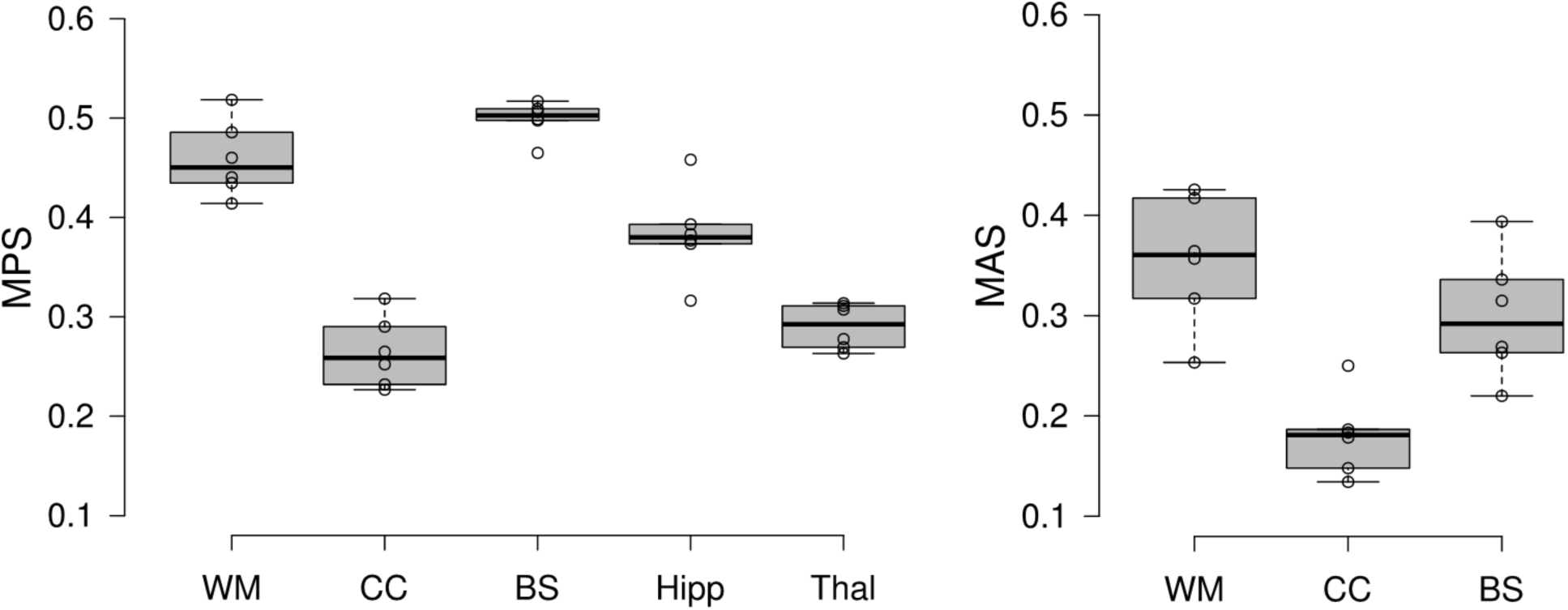
Boxplots of MPS and MAS at brain regions for the six head models. MAS shows a more extensive spread than MPS, indicating a larger inter-subject variability. The boxplots show the median, minimum, and maximum value.

### 3.6 Brain shape influence

The 50^th^ percentile female and 5^th^ percentile male have very close ICV, and brain volumes are also similar (5% difference). Thus, it would be interesting to analyze further to understand how brain strains vary between subjects with similar ICV but different shapes. Simulation results from the two subjects are compared, showing MPS in the cerebral cortex differs by 14.2% (**Fig. 12**a), occurring at a quite different location as shown in **Fig. 8**c, despite a similar pattern in strain distribution (**Fig. 12**b). For CC, not only the MPS value differs between the two, more importantly, the occurring time (**Fig. 12**c), and consequently, the strain distribution pattern when MPS occurs (**Fig. 12**d). MPS is located at the mid-body of CC in the 5^th^ percentile male and located at the anterior in the 50^th^ percentile female. The results provide evidence that for heads with similar ICV, the magnitude of MPS, location, and strain pattern can vary significantly due to head shape difference.

**Fig. 12.**
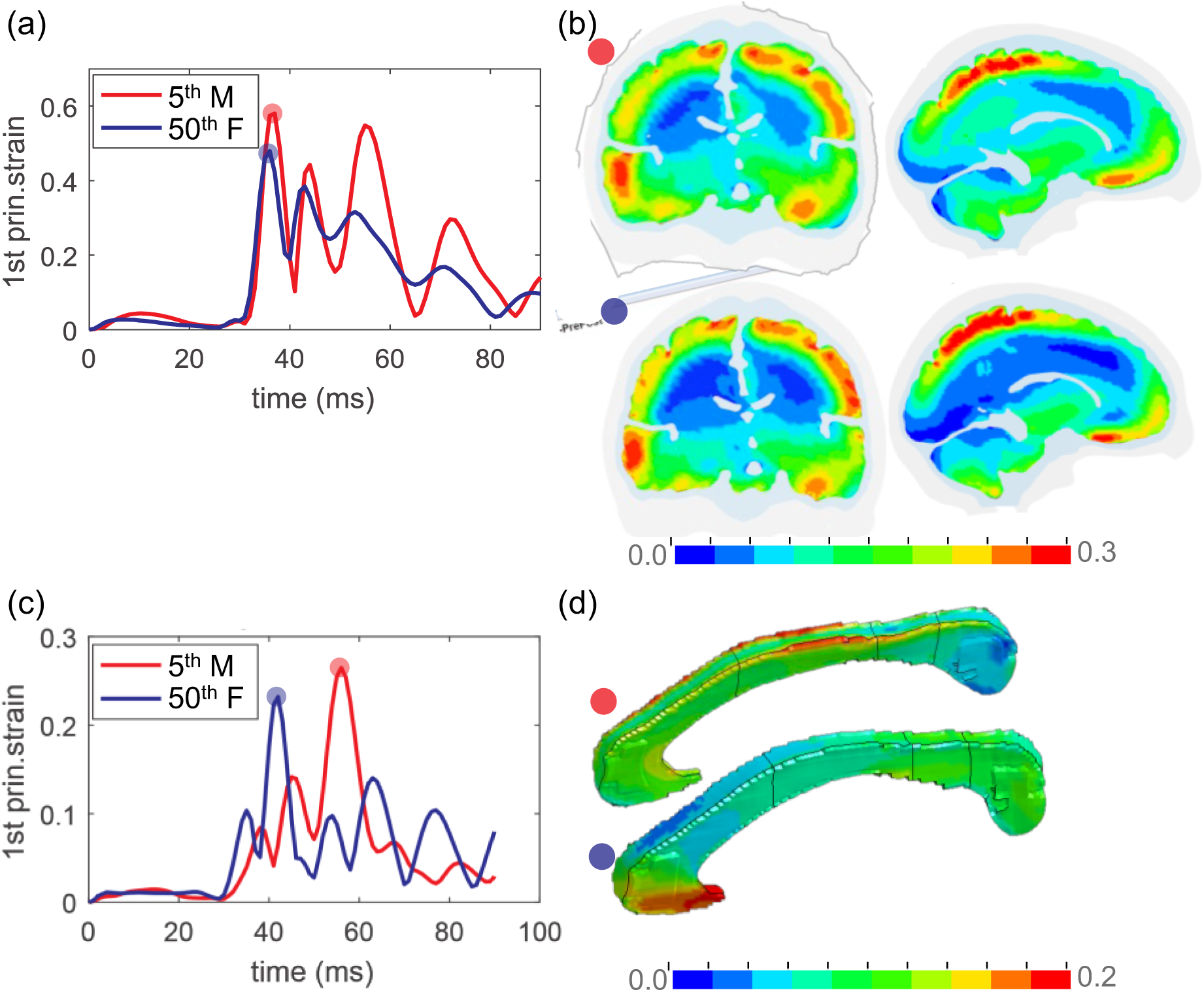
Brain strain response in the two subjects with almost the same ICV. (a) The time-history curve of MPS in the cerebral cortex. (b) Coronal and sagittal cross-sections of the 1^st^ principal G-L strain captured at MPS occurring time in both subjects. (c) The time-history curve of MPS in CC. (d) 1^st^ principal G-L strain distribution captured at the MPS occurring time in the respective subject.

## 4 Discussion

This study presents an anatomically detailed and personalizable head model with WM tracts embedded (the ADAPT head model), and a hierarchical image registration pipeline for subject-specific head model generation by mesh morphing. The model is validated against experimental brain motion and brain strain close to or at injury level, as well as intracranial pressure, showing overall CORA score comparable or higher than earlier models. The developed pipeline allows efficient generation of anatomically detailed subject-specific head models with satisfactory element quality. Subject-specific head models generated using this approach are shown to capture well the subjects’ head geometry for the six subjects of largely varying ICVs, both on a global level (cranial mask and the brain), and local brain regions as well as lateral ventricles. Brain strains of MPS and MAS show significant differences among the six subjects due to head/brain size & WM morphology variability, motivating the necessity for using subject-specific head models for evaluating brain injuries.

### 4.1 Head size influence on brain strain response

The significant difference in both MPS and MAS magnitude and locations among the subjects (**Fig. 8** - **Fig. 11**) seems to suggest ICV as a dominant factor influencing brain strain. Further, heads with larger ICV tend to have increased brain strains under the same loading, though not necessarily follow a monotonic increasing trend. Notably, the two subjects with very similar ICV show quite different strain patterns (**Fig. 12**), indicating head/brain shape may also be an important factor influencing brain strain response. It will be interesting to investigate in the future to clarify whether size or shape is more important than the other influencing brain mechanical response. For example, by using principal component analysis (PCA) (Wold et al. 1987) with more brain images, which may also allow identifying the characteristics of brain shape and WM morphologies that are most vulnerable to impact.

Percentile strain (e.g., 95^th^ percentile MPS) have been used in previous studies both for adults (Miller et al. 2019; Panzer et al. 2012; Wu et al. 2020; Wu et al. 2020; Beckwith et al. 2018) and child head models (Li and Kleiven 2018) to avoid potential numerical issues (e.g., strain concentration). This study uses MPS (i.e., 100^th^ percentile MPS) to compare between subjects. Using MPS also allows identifying the element with maximum strain and plotting its time-history curves, as presented in **Fig. 8**a, which is from a brain element on the cortical surface. Visual check shows the strain in the plotted element has no abrupt difference comparing with its neighboring elements at the cerebral cortex (i.e., sulci/gyri) attributed to the conforming mesh, giving some confidence in the plotted curves, as well as the identified varying locations among subjects (**Fig. 8**c). **Fig. 9** with strain curves plotted from all elements at brain regions also shows no curves jumping outside, brings further confidence on the reported MPS. Finally, the time-history strain curves shown in **Fig. 10**b from one element in the CC, despite the non-smooth interface between CC and neighboring brain elements due to the same brain material used, no abrupt strain differences are observed in the plotted element comparing with its neighboring elements.

Influences of head/brain size on brain mechanical response have been studied in the past. In a 2D numerical study by Prange et al. (1999), coronal rotational accelerations were applied to evaluate brain strain response due to brain size differences between adults and children. Kleiven and von Holst (2002) by globally scaling a 3D adult FE head model to six heads of varying dimensions, showed brain response increased almost monotonically from the smallest to the largest head under a linear acceleration. Similarly, as revealed in this current study, a larger ICV (relating to larger brain mass) tends to have a larger strain under the same impact, which is also suggested by Holbourn’s scaling principle (Ommaya et al. 1967), and more recent work by Wu et al. (2020) and Panzer et al. (2014). Of interest would be to investigate whether brain strains predicted from FE models follow or can be predicted by the acceleration-mass scaling law, which indeed has been studied by Prange et al. (1999). Their results demonstrated that the mass scaling relationship was not sufficient to produce brain strain distributions. Similarly, the models in the current study incorporating sulci and gyri, poses even greater challenges for such scaling laws to relate MPS with brain masses. The anatomically detailed subject-specific head models representing individual brain structural differences appear to be critical for revealing the new insights on brain size/shape influence (**Fig. 8** and **Fig. 12)** comparing with models that have smooth brain surface and by global scaling (Kleiven and von Holst 2002).

### 4.2 Image registration pipeline for mesh morphing

Image registration is a well-developed research area in the neuroimaging field, with algorithms ranging from global (e.g., rigid align, affine) to deformable registration, which allows obtaining dense displacement field reflecting the vast difference among subjects. However, most algorithms are developed with intended use within the neuroimaging field, e.g., for template generation, template guided segmentation, quantity group difference by registering subjects’ images to the same template (Oliveira and Tavares 2014; Toga and Thompson 2001). When applying for mesh morphing, higher requirement is imposed on the smoothness of the resulting displacement field than usually required in neuroimaging field, not only associated with FE models’ runnability but also prediction accuracy. For the developed hierarchical registration pipeline in this study, both the choice of Demons (Vercauteren et al. 2009) and Dramms algorithms (Ou et al. 2011), as well as the hierarchical design are essential to obtain displacements fields that allow generating subject-specific head models with competitive personalization accuracy, meanwhile with satisfactory element quality without mesh repairing.

Demons registration allows morphing brains with large differences; however, it tends to result in displacement fields that may lead to excessive element distortion according to our previous experience (Li et al. 2013; von Holst and Li 2013; von Holst et al. 2012). The hierarchical design of the pipeline is to utilize Demons’ capacity for handling large shape differences by performing Demons registration as a first step with binary cranial masks as input that allow obtaining a displacement field reflecting well overall cranial shape (mean DICE of 0.975, **Fig. 7**). Dramms registration is performed in the 2^nd^ step on the skull stripped T1W images inherited from the 1^st^ step. The focus of the 2^nd^ step registration is to align local brain structures and CSF (outer CSF and ventricles) by using brain MRI information without need of segmentation. The choice of Dramms is based on its promising performance due to its advancing in hierarchical attribute matching mutual-saliency mechanism (Ou et al. 2011). It will be interesting to investigate if other nonlinear algorithms would achieve good performance for the detailed brain or not, such as B-spline and Burr’s elastic previously used for morphing head models with smooth brains (Ji et al. 2015; Wu et al. 2019). Besides, other popular algorithms developed in the neuroimaging field (e.g., DRATEL by Ashburner (2007), and more found in Ou et al. (2014)) when used alone or replacing Dramms in the 2^nd^ step may result in better performance or not. Nevertheless, the proposed hierarchical registration pipeline leads to competitive performance in generating detailed subject-specific head models with satisfactory element quality without the need for further element repairing, meanwhile achieves DICE values comparable to or higher than that in the neuroimaging field (Ou et al. 2014).

With the established pipeline, new advanced registration algorithms developed in the neuroimaging field can be readily implemented to generate subject-specific models reflecting even better the intersubject-variability for both brain and WM fiber morphologies. The hierarchical image registration pipeline is not only applicable for this current ADAPT head model but also can be used to morph other head models as baseline, such as smoothed voxel head models. Further, due to its capacity for handling highly nonlinear warping, the pipeline can also be used to generate models with pathologies with brain structural changes such as decompressive craniotomy when the brain is expanding outside the skull (von Holst et al. 2012).

The proposed pipeline allows good alignment for local brain regions such as CC, achieving almost perfect alignment (DICE of 0.95) for CC for the smallest female, though less ideal (DICE of 0.85) for the largest female (Fig. 3). Note that the less ideal DICE values for some cases are not to be seen as performance indicator for the Dramms algorithm per se. Since the performance replies on input image alignment for Dramms, which can be tuned to achieve a better performance. Nevertheless, the DICE values for GM and WM are difficult to improve by registration. This issue could be improved by regrouping WM according to the registered neuroimages which will allow representing the subject’s WM accurately. Therefore, combining regrouping would allow the mesh morphing approach as an efficient approach for generating subject-specific models with competitive accuracy comparing with developing a model from scratch. For example, the Hexotic approach as described for the baseline model involves accurate image segmentation, generating surfaces, manual generation of membrane elements, which is a tedious process; sometimes maybe impossible due to the low quality of medical images for the subject of interest.

### 4.3 Brain-skull relative motion validation performance comparing with previous models

Regarding model validation performance on brain-skull relative motion, the ADAPT head model shows consistently higher CORA scores comparing with the original KTH head model, while has comparable CORA scores with the KTH-FSI model and the KTH detailed head model. The seven cases for brain-skull relative motion validation selected here are because they consistently have a duration longer than 40 ms. Besides, only reliable NDTs are included in CORA score calculation, justifying the same weight factor used for all NDTs (i.e., CORA score for each case is a mean of NDT curves). CORA scores from previous models are either newly calculated or recalculated in this study to ensure all values are directly comparable among models. In fact, CORA scores for the original KTH head model for C288-T3, C380-T4, C380-T6, C393-T3 have been presented earlier, showing comparable CORA scores with THUMS, GHBMC, see Table 8 in the study by Giordano and Kleiven (2016). Note despite this study followed the same CORA calculation method (with corridor method excluded) and used recommended global settings same as proposed by Giordano and Kleiven (2016), the same weighting factor is used for all the evaluated NDTs instead of using the proposed weights for each NDTs as listed in Table 5 (Giordano and Kleiven 2016). The reason is that seems no consensus has been reached among the research community on the proposed weighting factors and using equal weight for all NDTs is justified by only include reliable NDTs as plotted to allow easier comparison for future studies. Nevertheless, the CORA scores for the overlapping cases are close between the two studies, e.g., for C380-T4 5.26 shown in **Table 8** in Giordano and Kleiven (2016), close to 0.551 reported here (note differing by a factor of 10 in the calculation).

### 4.4 Experimental brain strain for head model validation

Regarding experimental b rain strain data used for model validation, the cluster brain strain presented in Zhou et al. (2019b) is used, which is recalculated based on the original brain-skull relative motion experimental data from Hardy et al. (2007) using a tetra approach instead of a triad approach used in the original study (Hardy et al. 2007). Though it’s well recognized (Zou et al. 2007; Zhao and Ji 2020) and recently has been extensively verified (Zhou et al. 2019b; Zhou et al. 2018) that a model validated against brain-skull relative motion may not necessarily guarantee its strain prediction accuracy. Therefore, it’s suggested that a head model with an intended use for strain prediction should be validated against experimental brain strain data. However, despite the availability of the strain data presented in Hardy et al. (2007) along with the brain motion data, it’s seldomly used for head model validation in contrast to motion data being widely used. This may be partially attributed to concerns on the quality of strain data, especially the initially reported ∼ 2–5% peak strains are rather low for an injurious impact (see a detailed discussion by Zhao and Ji (2020) and references therein).

To address the quality concerns on strain data originally presented in Hardy et al. (2007), a two-step effort has been undertaken recently. As a first step, Zhou et al. (2018) reanalyzed and updated the brain strain data using the same triad approach and developed three criteria to assess the eligibility of the NDT clusters suitable for strain calculation. As a second step, a tetra approach is used for brain strain calculation (Zhou et al. 2019b), reflecting better the 3D experimental brain deformation than the triad approach. Note the tetra or triad approach is just one of another way to estimate brain strain from NDT motion, alternative approach has been proposed also, e.g., a generalized marker-based strain sampling approach to estimate and compare regional strains (Zhao and Ji 2020). Indeed, experimental brain strain data calculated by the tetra approach are much larger than by the triad approach, indicating the original triad strains (Hardy et al. 2007), also the reanalyzed triad strains (Zhou et al. 2018) largely underestimated the experimental brain strain. Thus, the experimental brain strain data calculated by the triad approach is not recommended to be used for head model validation due to its large underestimation of the real brain strain in the experiment.

Note that for nearly incompressible material as the brain, shear strains should be close to principal strains, indeed as the ADAPT model predicted for all the seven clusters (see **Fig. 6** simulated shear strain and principal strain). However, unexpected large differences are observed between experimental principal strain and experimental shear strain (**Fig. 6)**. However according to a study by Bradshaw (2003), it is quantified that for brain tissue, the principal strain should fall within the range of 2/3 to 4/3 of the maximum shear strain based on the theoretical interpretation. While when reanalyze the experimental strain data from Zhou et al. (2019b), of all the 15 clusters, the experimental principal strain consistently falls within the shear strain band, correlating with the theoretical finding by Bradshaw (2003). Such correlation seems to indicate an acceptable quality of the 15 cluster brain strain dataset including the 7 used here for validation. Therefore, the recalculated cluster brain strains using the tetra approach have been verified and justified thoroughly (Zhou et al. 2019b), and can be taken as experimental brain strain for validating brain strain response of FE head models.

### 4.5 Brain strain validation performance

Seven cluster strains are chosen (out of 15 in total) for model validation as they consistently have duration over 40 ms. The shape and phase show a good match between ADAPT model simulated and experimental brain strain, but a large discrepancy in magnitude is observed in most clusters (**Fig. 6**). Further, the simulated brain strains are consistently lower than the experimental strain in all the evaluated clusters except for C380-T1 C1. It will be interesting to study more clusters, especially, experimental strains for C241-T5 C2, C241-T6 C2, C393-T2 C1 have much lower peak strains (all <0.1) (Zhou et al. 2019b). It’s expected that for some of these clusters, simulated brain strain may be larger than experimental strain, which, however, is yet to be performed. This also reminds one important difference comparing with NDT motion data (many NDT curves even for one case, e.g. 14 NDTs and each has XYZ displacements), that choosing only several or certain cluster strains may lead to a biased impression that the simulated brain strain too low compared with experimental data; this further reminds that caution should be taken when tuning material parameters to satisfy brain strain performance when only use a limited number of clusters (as shown in **Fig. 6** for the seven clusters selected in this study consistently shows lower strain in the model, but may not be the case when more cluster are evaluated as discussed above).

Therefore, the large peak discrepancy between ADAPT and experimental cluster strain is suggested not to be seen as a concern for the model performance, rather it opens a question not specific to this model: How to use the experimental cluster strain for validation for the models that chose to use this strain data. Especially, how to evaluate model performance when only a few cluster curves are available versus NDT motion curves where are many. Further, how to extract brain strain for a model with coarse mesh. Note CORA scores for brain strain is actually higher than brain motion despite large magnitude discrepancy, seems to remind CORA scores may not be a proper index, which thus raises a question on how to weight between NDT motion and strain to reach an overall biofidelity ranking for a model.

The recalculated experimental cluster brain strain data is a result of the two-step efforts (Zhou et al. 2018, Zhou et al. 2019b) attempted to best utilize the NDT motions close to or at injury level originally measured by Hardy et al. (2007). The recalculated experimental cluster brain strain data provides a possibility to evaluate head models’ brain strain prediction capacity; however, there are challenges, as mentioned above, that need integrated effort among the research community to allow its proper use for head model validation. Efforts toward this direction has been initiated (Zhao and Ji, 2020), and a thorough discussion on the use of strain data for model validation and the insufficiency of CORA scores for model biofidelity rating can be found therein.

### 4.6 Material modeling choice for the brain, falx, tentorium, dura and pia

The ADAPT model uses the same material properties for the brain as the previously validated KTH head model (Kleiven 2007) with a coarse brain mesh. Indeed, studies have demonstrated that models with finer meshes would lead to the prediction of larger brain-skull relative motions and larger brain strains with the same set of material properties (Giudice et al. 2019; Zhao and Ji 2019). It is thus suggested that it may not be appropriate to adopt material properties of the brain obtained from a coarse mesh to a model with a much finer mesh, and vice versa (Zhao and Ji 2019). This is indeed a valid concern especially if brain parameters were obtained by adjusting/optimization to satisfy model validation, resulting model-specific material properties that may not be directly translatable to other models, especially with largely different brain mesh sizes. Keeping this in mind, it seems questionable to adopt brain material properties from the KTH head model to ADAPT. However, note that the brain material properties presented in (Kleiven 2007) were based on careful analysis and data fitting of experimental data, thus can be considered as model-independent and translatable to the ADAPT model. While updating falx/tentorium material properties with a nonlinear model instead of using the same linear elastic model (Young’s modulus of 31.5 MPa) is based findings from a recent study (Ho et al. 2017), showing the importance of using nonlinear material models that allow reflecting the nonlinear properties falx/tentorium as shown from experiments. Similarly, pia mater is updated in the ADAPT model using a nonlinear model based on experimental data. Due to significant differences in mesh sizes, updated material model for pia/falx/tentorium/dura, as well as sliding contact in the KTH head model versus continuous mesh in ADAPT, different brain responses are expected from the two models under the same loadings. Indeed, ADAPT model, in general, has a larger brain-skull relative motion for most NDTs (see **Fig. A3** to **Fig. A5**), and has higher CORA scores for the seven cases than the KTH head model (**Table 3**). Further, the detailed brain morphology (sulci/gyri) included in the current model may potentially alter brain tissue response, considering the CSF penetrates and pia mater holding the brain at the sulci/gyri comparing with models with smooth brain surface. Nevertheless, a more systematic investigation is required to study the potential prediction differences between ADAPT and the original KTH head model. Consequently, injury criteria developed (Kleiven 2007) may not be directly applicable to the ADAPT model, and care should be taken when using the ADAPT model for injury prediction based on existing injury criteria developed from the KTH head model and other head models.

Mechanically anisotropic brain tissue models, in particular, the Gasser, Ogden and Holzapfel (GOH) model have been implemented in FE head models (Giordano et al. 2014; Giordano and Kleiven 2014b; Giordano et al. 2017; Zhao and Ji 2018), by connecting anisotropic diffusion properties from DTI to mechanical anisotropy (Giordano and Kleiven 2014a). Some experimental studies suggest that brain tissue is mechanically isotropic, e.g., Budday et al. (2017), while others suggest brain tissue show significant directional trends, e.g., Prange and Margulies (2002), Feng et al. (2017). More discussion on the controversies can be found in a review study by Budday et al. (2019). Nevertheless, both isotropic and anisotropic models are being used in head models for studying TBIs; see review studies (Giudice et al. 2019; Madhukar and Ostoja-Starzewski 2019). In the current study, isotropic hyperviscoelastic material model is used for brain tissue, and axonal strains are calculated by projecting strain tensors to WM fiber tract directions as earlier studies have shown MAS as a potentially improved predictor for brain injury (Sullivan et al. 2015; Zhao et al. 2017; Zhao et al. 2016) using the same approach or explicitly model brain as GOH (Giordano and Kleiven 2014b). Nevertheless, how brain material difference (e.g., isotropic versus anisotropic) may influence brain strain in an anatomically detailed head model is yet to be studied.

### 4.7 Limitation and future work

There are some limitations in the ADAPT model to be noted. The ADAPT model has conforming mesh at all interfaces between the entire brain and CSF (i.e., outer CSF-brain/dura, ventricle-brain). Still, non-smooth boundaries exist at brain subregions, including GM and WM interface. The subregions of the brain, and diploe porous skull bone are grouped according to the spatial correspondence with the segmented images via an automatic script, which seems often similarly practiced in other head models as well, e.g., WM meshes manually picked according to geometry data in the GHBMC (Mao et al. 2013), and the same for the KTH head model (Kleiven 2007). Despite the non-smoothness, strain concentration is often not a concern, especially if using the same material properties for the entire brain as done in this study. Further, a continuous mesh is used throughout the ADAPT model, which can be further improved by including FSI at the brain-ventricle interface as done earlier (Zhou et al. 2020), and further to implement FSI at the brain-skull interface (Zhou et al. 2019a) in the future, despite a technical challenge to implement FSI on the complex sulci and gyri than a smooth brain model. Further, the model doesn’t validate well with the chosen *in vivo* data, and future investigations are needed to improve the model’s capacity for predicting brain response under non-injurious low impact loading. The ADAPT model also has substantially longer simulation runtime compared with models with fewer elements. Finally, the results presented in Sections 3.4, 3.5, 3.6 are to be seen as parametrical studies based on a validated baseline model, highlighting the differences in brain strain prediction among individuals under the same impact loading condition.

## 5 Conclusion

This study presents the development of an anatomically detailed head model with conforming hexahedral mesh (the ADAPT head model) equipped with a hierarchical image registration pipeline for efficient generation of subject-specific models by mesh morphing. The model is validated against brain-skull relative, brain strain, and intracranial pressure, showing comparable performance with previous models. The six-subject specific head models generated using the ADAPT model and the pipeline demonstrate the capacity of the ADAPT model and the pipeline for fast generation of anatomically detailed subject-specific head models with largely varying brain sizes/shapes with competitive personalization accuracy on capturing individual’s brain structures. The simulation results show significant differences in brain strain and axonal strain, motivating the necessity for using subject-specific head models for evaluating brain injuries. The ADAPT model due to its uniqueness of the complete ventricular systems, including 3^rd^ and 4^th^ ventricles in connection with outer CSF via aqueduct, together with conforming meshed sulci and gyri and subject-specific WM fibers, could potentially provide new insights into TBI mechanisms. The verified performance of the ADAPT head model equipped with the personalization approach addresses the challenges in subject-specific FE model generation, opening an opportunity for studying for studying personalized brain responses, as well as developing personalized head protection systems. The research community may find the hierarchical registration pipeline useful to morph other anatomically detailed head models, such as smoothed-voxel head models.

## Acknowledgments

The authors would like to acknowledge the earlier version of the Python code from Chiara Giordano, which was adapted and used in this study. Our special thanks go to Dr. Yangming Ou and Dr. Steve Pieper, who generously spent their time advising Dramms registration, and to Dr. Loïc Maréchal who provided great support for the usage of the software Hexotic. The authors thank the three anonymous reviewers and Prof. Peter J. Hunter for their stimulating comments and valuable suggestions that substantially improved this paper. The WUM HCP data of the six subjects used in this study were provided [in part] by the Human Connectome Project, WU-Minn Consortium (Principal Investigators: David Van Essen and Kamil Ugurbil; 1U54MH091657) funded by the 16 NIH Institutes and Centers that support the NIH Blueprint for Neuroscience Research; and by the McDonnell Center for Systems Neuroscience at Washington University. The computations were performed on resources provided by SNIC through Center for High Performance Computing (PDC) at KTH under Project SNIC 2019/3-509 and SNIC 2019/3-409. Gong Jing at KTH PDC center is acknowledged for assistance for the installation of needed software on the PDC resources. The present study was supported by research funds from KTH-Royal Institute of Technology, Stockholm, the Swedish Research Council Grants (nr. 2016-04203 and nr. 2016-05314), and European Union’s Horizon 2020 research and innovation programme under the Marie-Curie Grant Agreement No. 642662.

## Electronic Supplementary Material

### 1. Supplementary Validation Results of the Head Model

Validation setup and additional validation results for the ADAPT head model.

### 2. Supplementary Videos

Brain tissue response of the 1^st^ principal strain during the simulated entire concussion impact for two subjects: the smallest (female) and largest (male).

## Appendix 1 Personalization accuracy and mesh quality of the personalized models by morphing

The baseline T1W image **Fig. A1**a) is morphed to a personalized T1W image for each subject (**Fig. A1**b) using the procedure presented in Sec. 2.4. The personalized T1W image is paired with the subject-specific mesh. Thus a comparison between it and the subject’s own T1W (**Fig. A1**c) (the ground truth) allows evaluating how the subject-specific model reflects the subject’s head geometry. The results show not only the overall head shape, but also internal brain structures are well correlated (**Fig. A1** b, c). The personalized T1W images (**Fig. A1**b) and subject T1W (**Fig. A1**c) are then segmented for the cranial mask, brain, and seven local brain regions illustrated in **Fig. 4** in the main text for DICE calculation.

**Fig. A1.**
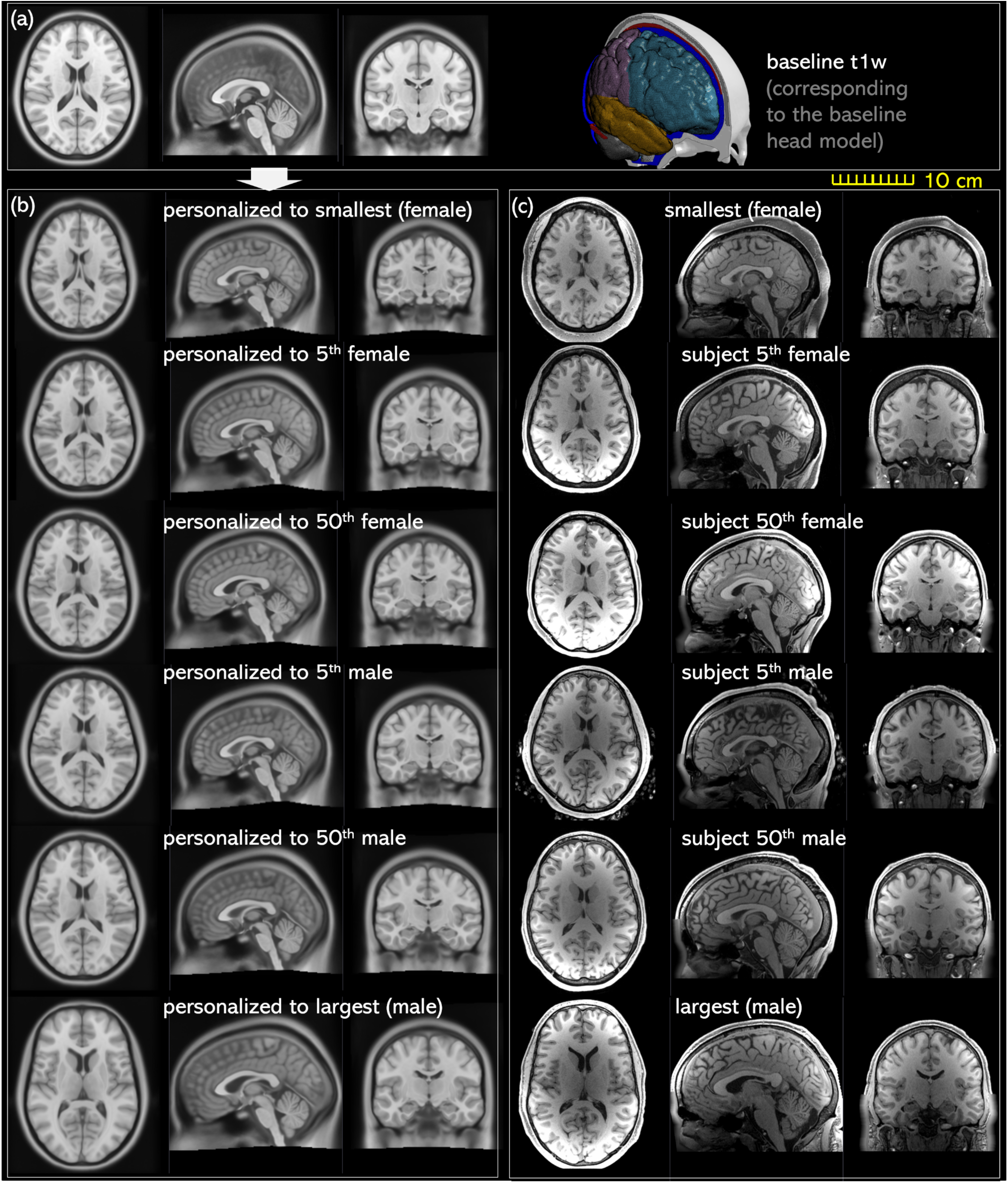
Baseline T1W image (a) and personalized T1W image (left column) comparing with the ground truth of the T1W image of the subject. Transverse, sagittal, and coronal cross-sections are captured for each image.

DICE coefficient for the six subjects corresponding to the boxplot **Fig. 7** in the main text.

**Table A1.**
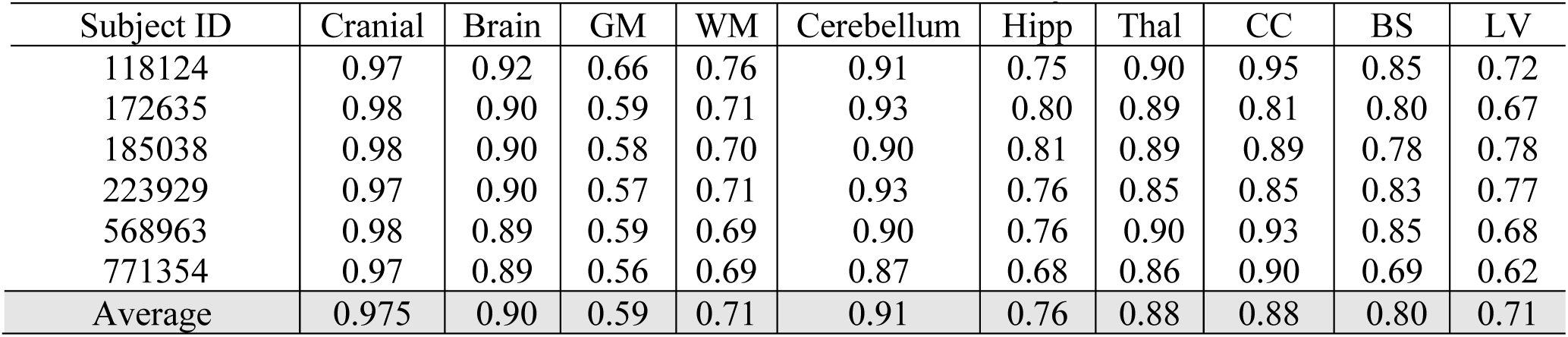
DICE coefficients for the six subjects.

**Fig. A2.**
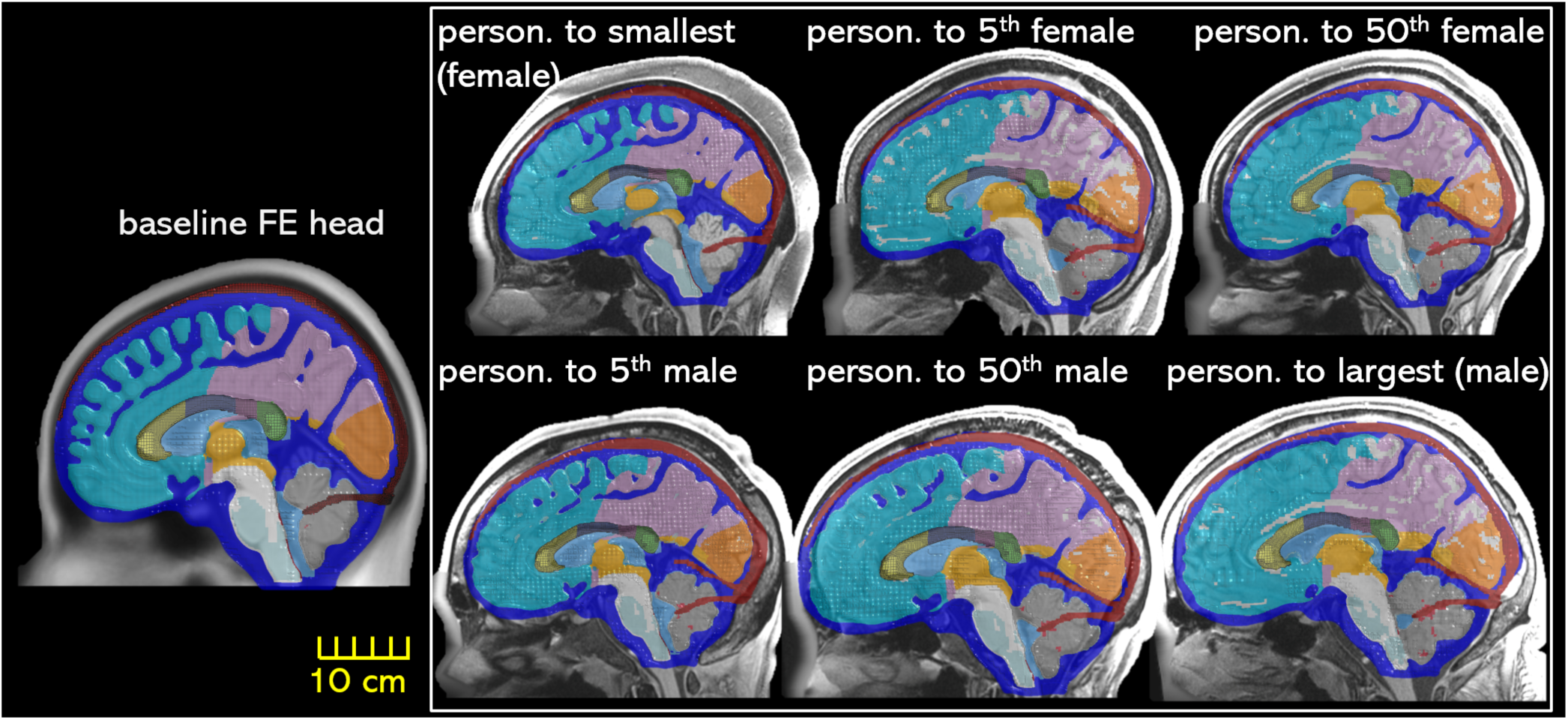
FE mesh of the six generated subject-specific head models by morphing. Sagittal section of the six head FE mesh is overplayed with the subject’s T1W image. The mesh is made half transparent to show the correspondence between the mesh and the ground truth T1W image.

**Table A2.** Element quality of the ADAPT model and the six subject-specific head models generated by morphing.

| Head model | Element quality Index |  |  |  |  |  |  |  |  |  |  |  |
| --- | --- | --- | --- | --- | --- | --- | --- | --- | --- | --- | --- | --- |
|  | Jacobian<br>≥0.5 |  | Warpage (°)<br>≤30 |  | Skew (°)<br>≤60 |  | Aspect ratio<br>≤8 |  | Min. angle (°)<br>≥30 |  | Max angle (°)<br>≤150 |  |
|  | percent | min | percent | max | percent | max | percent | max | percent | min | percent | max |
| ADAPT | 98% | 0.45 | 92% | 111.76 | 99.9% | 69.95 | 99.9% | 6.62 | 99.8% | 17.98 | 99.9% | 161.94 |
| Personalized models |  |  |  |  |  |  |  |  |  |  |  |  |
| Smallest (female) | 95% | 0.27 | 91% | 134.24 | 99.9% | 70.75 | 99.9% | 7.83 | 99.9% | 13.65 | 98% | 168.00 |
| 5<sup>th</sup> perc. female | 95% | 0.28 | 91% | 117.11 | 99.9% | 70.61 | 99.9% | 8.64 | 99.9% | 11.65 | 98% | 168.70 |
| 50<sup>th</sup> perc. female | 94% | 0.21 | 91% | 121.79 | 99.9% | 74.48 | 99.9% | 7.79 | 99.0% | 12.24 | 98% | 203.35 |
| 5<sup>th</sup> perc. male | 95% | 0.32 | 91% | 125.85 | 99.9% | 69.22 | 99.9% | 9.05 | 99.9% | 14.57 | 98% | 168.33 |
| 50<sup>th</sup> perc. male | 95% | 0.26 | 92% | 114.11 | 99.9% | 70.40 | 99.9% | 8.20 | 99.9% | 11.54 | 98% | 170.14 |
| Largest (male) | 95% | 0.31 | 92% | 110.42 | 99.9% | 73.14 | 99.9% | 7.38 | 99.9% | 14.27 | 98% | 169.01 |

## Appendix 2 Brain-skull relative motion and brain strain validation comparing with previous head models

Brain-skull relative motion for three cases predicted from the ADAPT, and the original KTH head model (Kleiven 2007) are compared for C288-T3 (**Fig. A3**), C380-T1 (**Fig. A4**), C380-T2 (**Fig. A5**), together with experimental data from (Hardy et al. 2007). Note that C288-T3 NDT8 has a duration of 39 ms (<40 ms). Thus NDTs from 1 to 14 except NDT8 are included in the plot also in the CORA calculation. Besides, for the six impacts delivered to subject C380 (i.e., C380-T1 to T6), the motion of NDT9 failed to exhibit a motion trend toward its starting position (details found in (Zhou et al. 2018). Thus NDT1-14 except NDT9 are included in the plot also in the CORA calculation. Finally, for C393-T3, NDT13, therefore, NDT1-14 are included in the plot also in the CORA calculation.

CORA scores calculated for the KTH detailed head model (Zhou et al. 2019b) on brain strain are presented in **Table A3** in comparison with the ADAPT head model presented in **Table 4** in the main text.

**Table A3.** CORA scores of the KTH detailed head model (Zhou et al. 2019b) on principal and shear strain.

| Exp. cases<br>(Zhou et al. 2019b) | Principal strain |  |  |  | Shear strain |  |  |  |
| --- | --- | --- | --- | --- | --- | --- | --- | --- |
|  | V<br>(Shape) | G<br>(Size) | P<br>(Phase) | CORA score | V<br>(Shape) | G<br>(Size) | P<br>(Phase) | CORA<br>score |
| C288-T3 C1 | 0.929 | 0.202 | 0.797 | 0.643 | 0.954 | 0.229 | 0.919 | 0.701 |
| C380-T1 C1 | 0.981 | 0.843 | 0.797 | 0.874 | 0.975 | 0.656 | 0.919 | 0.850 |
| C380-T2 C1 | 0.993 | 0.286 | 0.999 | 0.760 | 0.986 | 0.409 | 0.999 | 0.798 |
| C380-T3 C1 | 0.982 | 0.239 | 0.999 | 0.740 | 0.982 | 0.301 | 0.919 | 0.734 |
| C380-T4 C1 | 0.950 | 0.218 | 0.919 | 0.696 | 0.962 | 0.258 | 0.797 | 0.672 |
| C380-T6 C1 | 0.971 | 0.204 | 0.685 | 0.620 | 0.968 | 0.281 | 0.685 | 0.645 |
| C393-T3 C1 | 0.942 | 0.994 | 0.804 | 0.913 | 0.935 | 0.671 | 0.804 | 0.803 |
| **Average** |  |  |  | **0.749±0.111** |  |  |  | **0.743±0.076** |

Notes: <sup>1</sup> CORA scores for the first six cases newly calculated based on curves reported in the original study (as CORA scores were not reported). For case C393-T3, simulation is for this study, with CORA newly calculated.

**Fig. A3.**
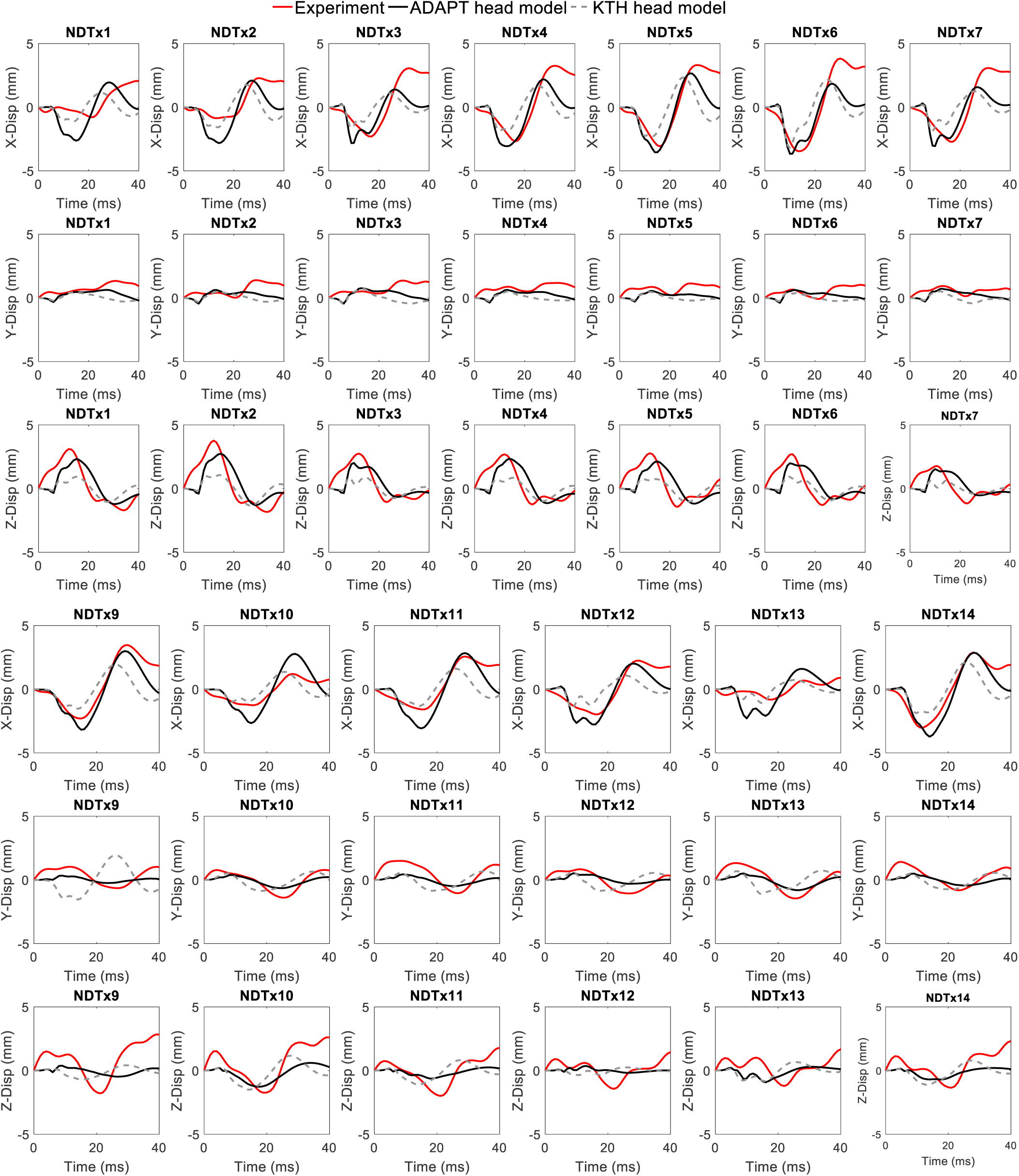
Comparison between experimental and simulated brain-skull relative motion by the ADAPT head model and KTH head model (Kleiven, 2007) for the experiment C288-T3 (Hardy et al. 2007).

**Fig. A4.**
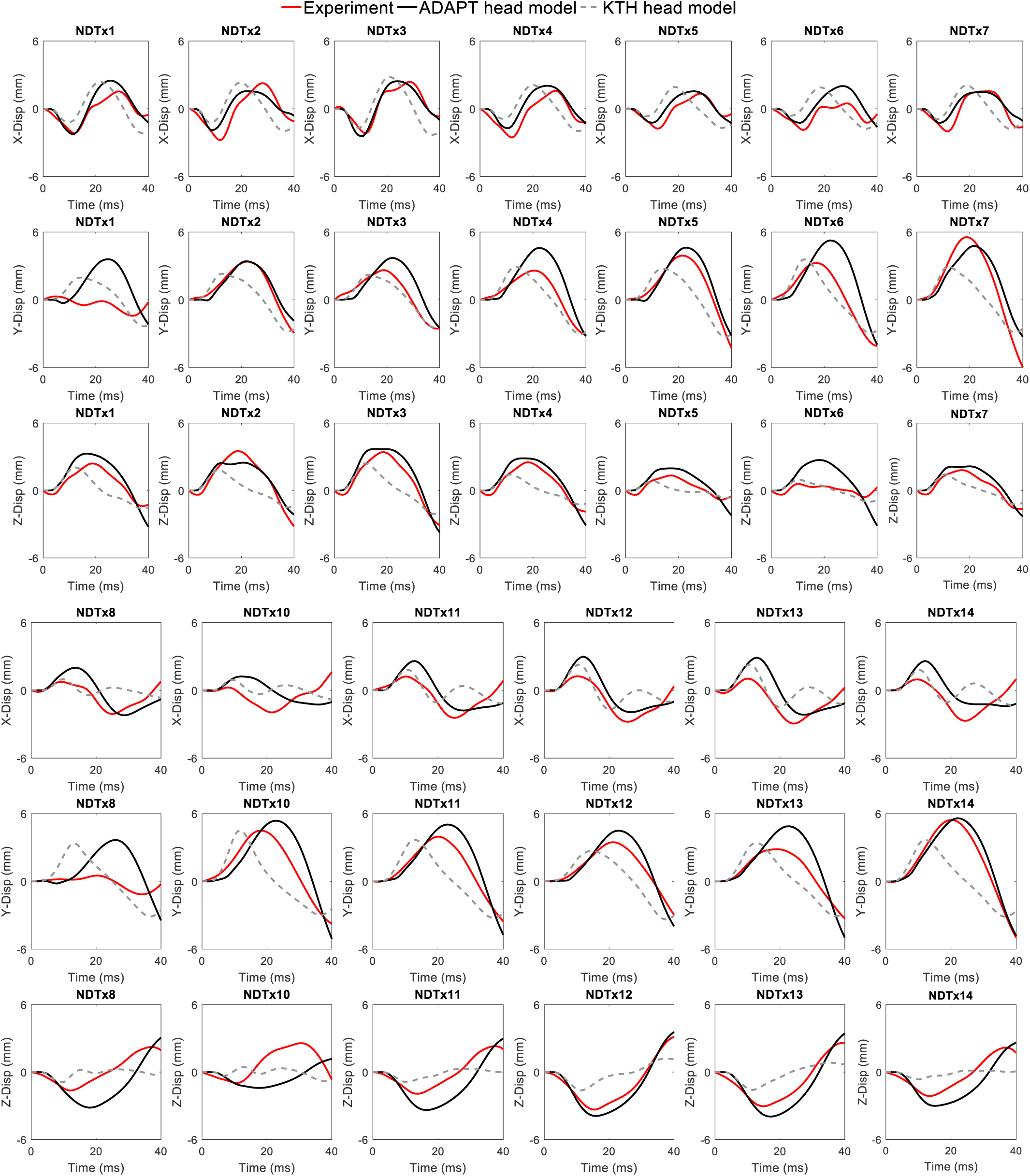
Comparison between experimental and simulated brain-skull relative motion by the ADAPT head model and KTH head model (Kleiven, 2007) for the experiment C380-T1 (Hardy et al. 2007).

**Fig. A5.**
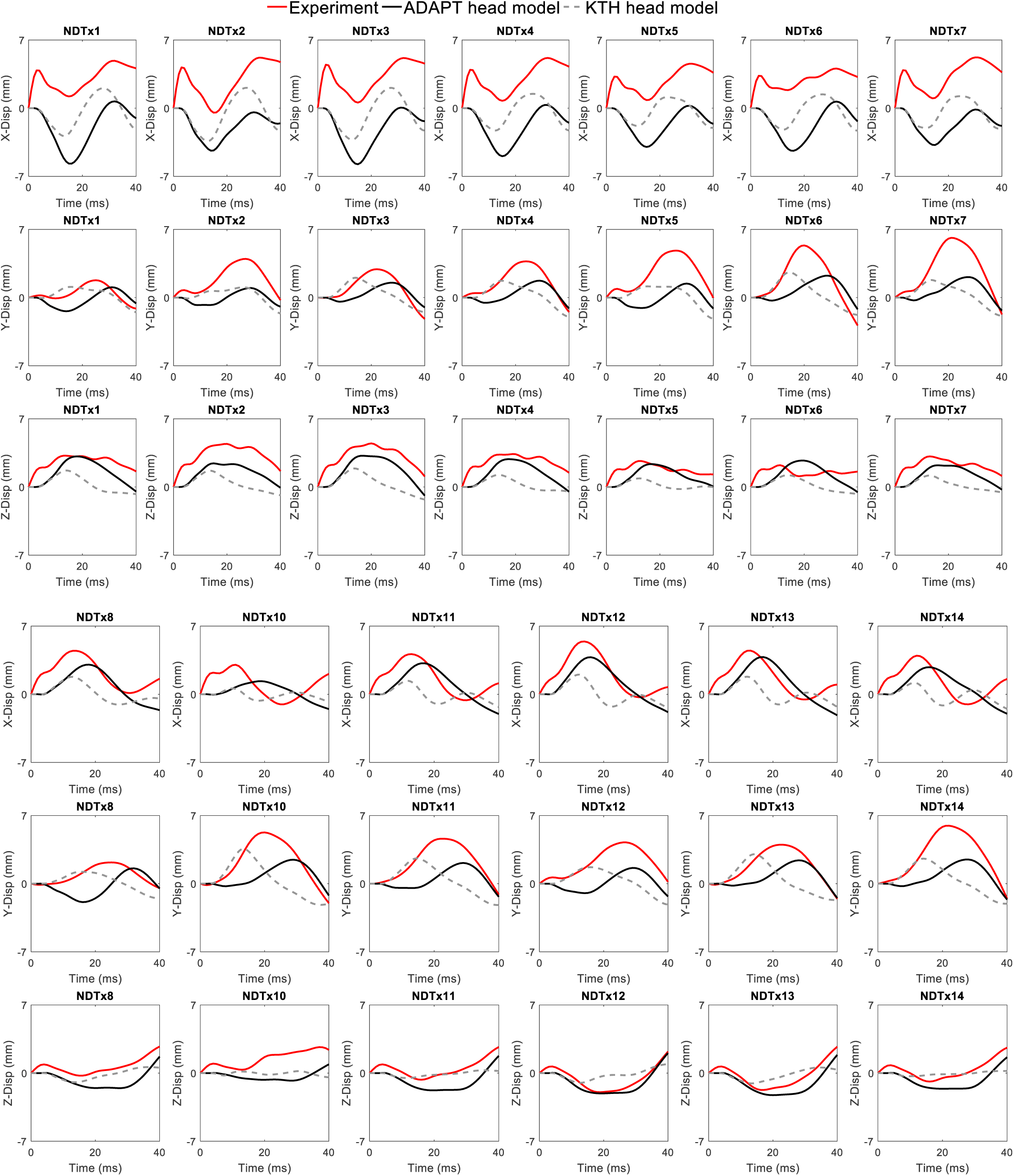
Comparison between experimental and simulated brain-skull relative motion by the ADAPT head model and KTH head model (Kleiven, 2007) for the experiment C380-T2 (Hardy et al. 2007).

## Appendix 3 Brain strains predicted from the six personalized models

The MPS and MAS for brain regions are presented in **Table A4** &**Table A5**, corresponding to the boxplot in **Fig. 11** in the main text.

**Table A4.**
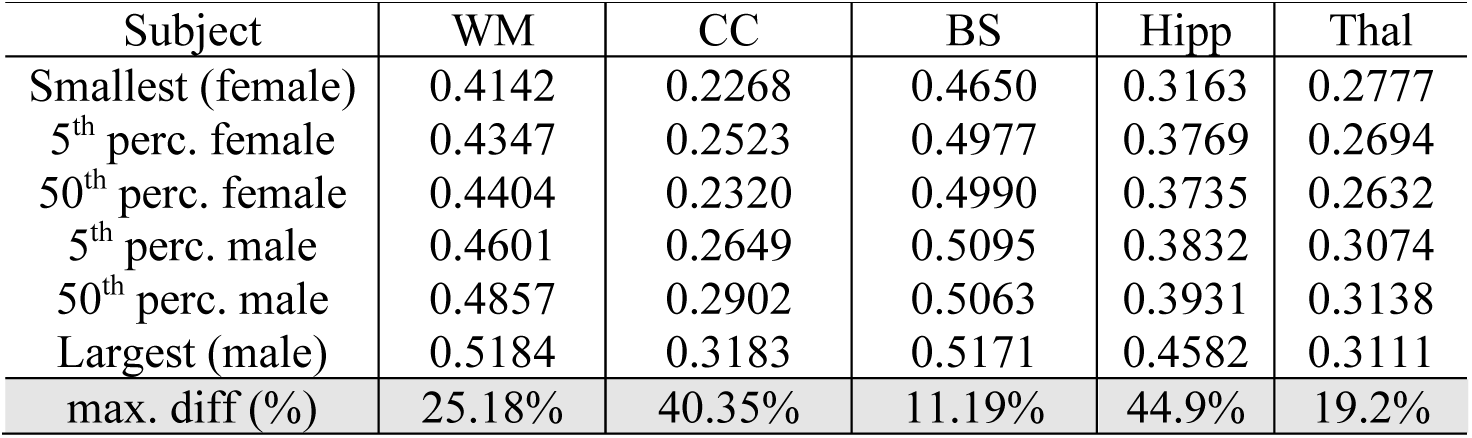
MPS for brain regions predicted from the six subject-specific head models.

**Table A5.**
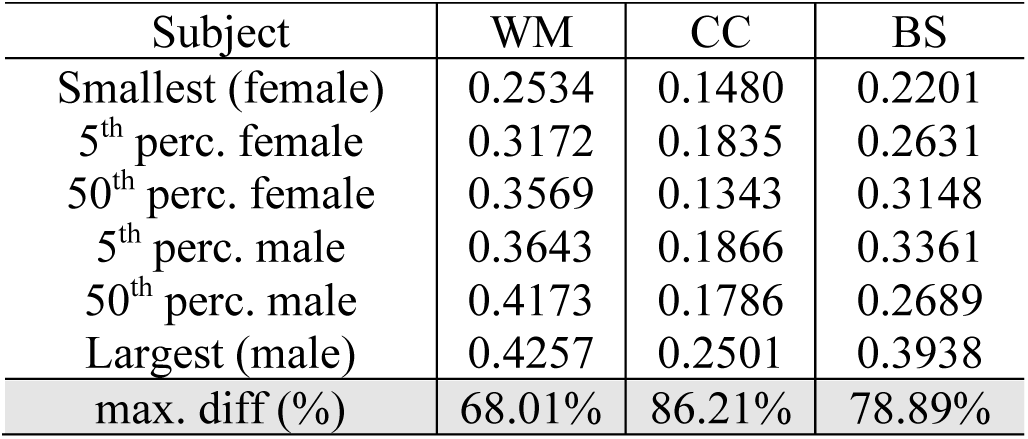
MAS for brain regions predicted from the six subject-specific head models.

