## Supplemental Material for "An anatomically accurate and personalizable head injury model: Significance of brain and white matter tract morphological variability on strain"

Supplementary Material Summary

#### 1. Supplementary Validation Results of the Head Model

Validation setup and additional validation results for the ADAPT head model.

#### 2. Supplementary Videos

Brain tissue response of the 1st principal strain during the simulated entire concussion impact for two subjects: the smallest (female) and largest (male).

### 1. Head model validation summary

The baseline head model is validated against experimental data of brain-skull relative motion in Hardy et al. (2007)[1] and the recalculated cluster brain strain[2]. To approximate the specimen anthropometry, the models are scaled independently in directions of both the depth and breadth to match the reported cadaveric head sizes. Intracranial pressure response of the FE head model is compared with recordings from experiment No. 37 conducted by Nahum et al. (1977)[3]. Further, to evaluate the model performance in predicting brain response in living subjects, brain-skull relative motion and brain strain are compared with the experimental displacements and strains measured in the human volunteer using tagged MRI during mild frontal impact from Feng et al. (2010)[4]. Details regarding model setup and results are presented in the following sections.

### 1. Validation setup

#### 1.1 Brain-skull relative motion & brain strain validation

For the brain-skull relative motion validation, seven representative cases with motion interval of embedded NDTs over 40-ms are selected, including one sagittal impact (C288-T3), one horizontal impact (C380-T2), and five coronal impacts (C380-T1, C380-T3, C380-T4, C380-T6, and C393-T3). Strain in cluster 1 (C1) of these 7 selected cases are further used for strain validation. To numerically replicate the experimental impacts, the recorded head kinematic curves are imposed to the node which locates at the center of gravity of the corresponding cadaveric head and is rigidly attached to the skull. To approximate the specimen anthropometry, the models are scaled independently in directions of both the depth and breadth to match the reported cadaveric head sizes. The node nearest to the start position of an experimental NDT target is taken as the marker location in the model. Motions of the identified nodes with respect to the skull along three anatomical coordinate directions are obtained from the whole head model simulation. Following the NDT tetra approach described[2], the initial positions of the identified nodes and the nodal motion responses predicted in the model are used to calculate the strain responses, specifically first principal Green-Lagrange strain and shear Green-Lagrange strain, of the brain model.

#### 1.2 Intracranial pressure compare with experimental data from Nahum et al. 1977

Intracranial pressure response of the FE head model is compared with recordings from experiment No. 37 conducted by Nahum et al. (1977)[3], where impacts to the forehead by a padded impactor were performed and pressure secondary to the delivered impact were measured at four sites: (1) frontal lobe adjacent to the impact contact area; (2) inferior to the lambdoidal suture in the occipital bone; (3) immediately posterior and superior to the coronal and squamosal suture in the parietal area; (4) posterior fossa in the occipital area. Following the experiment setting, the model is scaled to match the dimension of the corresponding cadaveric head and rotated 45° forward. A cylindrical impactor with padding materials at the impacting end is developed to deliver impact to the head model, similar to the strategy adopted earlier[5,6].

#### 1.3 Brain motion & strain compare with in-vivo volunteer experimental data from Feng et al. (2010)

Experimental measurements of subject 3 (S3) from Feng et al. (2010)[4]. are selected as validation targets here. To reproduce the experimental set-up with the FE head model, the estimated skull rigid-body motion is used as input to drive the motion of a node near the foramen magnum that is rigidly attached to the skull and defined as the skull origin. Three components of the skull motion are defined, including linear displacement in the anterior-posterior direction, linear displacement in the inferior-posterior direction and angular displacement among the axis perpendicular to the plane of motion.

Brain displacement with respect to the skull and brain strain in a 2D slice located in the left hemisphere at 1cm from the mid-sagittal plane (**Suppl. Fig. 6**) shows that numerically predicted brain displacement and strain are much smaller than the experimental measurements. Such results indicate that the ICBM head model cannot be used as a prediction tool based on brain displacement and strain under the voluntary loading scenarios.

#### 2 Validation results

Time-history curves of the brain-skull relative motion from the ADAPT head model and experimental data by Hardy et al. (2007)[1] are presented for C380-T3 (**Suppl. Fig. 1**), C380-T4 (**Suppl. Fig. 2**), C380-T6 (**Suppl. Fig. 3**), and C393-T3 (**Suppl. Fig. 4**), while the results for C288-T3, C380-T1 and C380-T2 are presented in the Appendix of this study, together with CORA scores presented in the main text of this study.

Pressure-time history curves of the cortical brain elements at the four aforementioned locations are compared with the measurements in **Suppl. Fig. 5**, with CORA scores presented in the main text of the study. Considering the obvious differences between the measured data at two occipital sites (Exp. Occ1 and Exp. Occ2 in **Suppl. Fig. 5**) and the unavailability of accurate measuring locations, CORA score for the pressure response at the occipital lobe is not calculated. It is worth mentioning that the model estimated pressure falls within the range defined by the experimental data, indicating the plausibility of pressure response at the occipital lobe.

Brain displacement with respect to the skull and brain strain in a 2D slice located in the left hemisphere at 1cm from the mid-sagittal plane is reported in **Suppl. Fig. 6**.

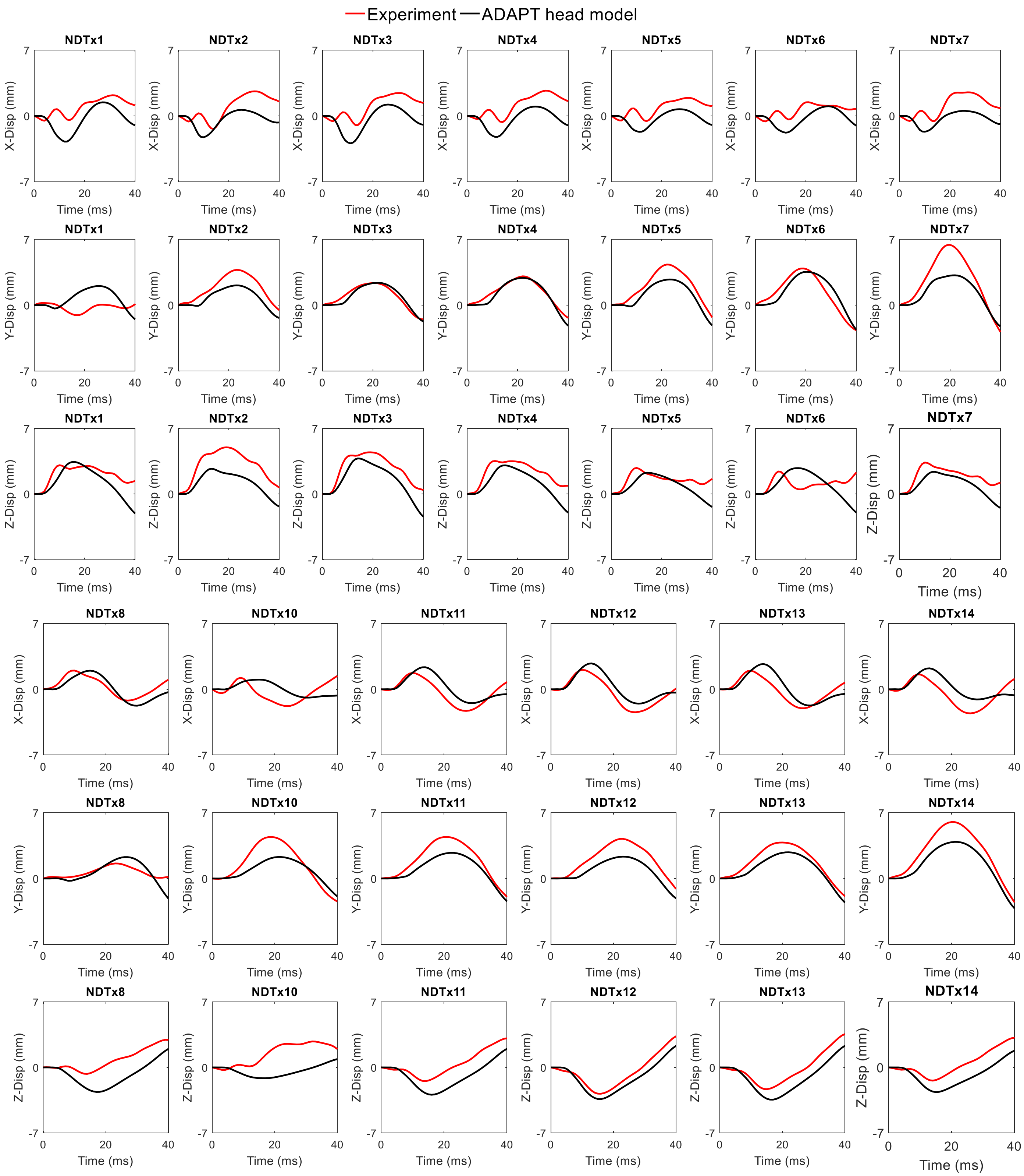


**Suppl. Fig. 1** Comparison between experimental motion and simulated brain-skull relative motion for the experiment C380-T3.

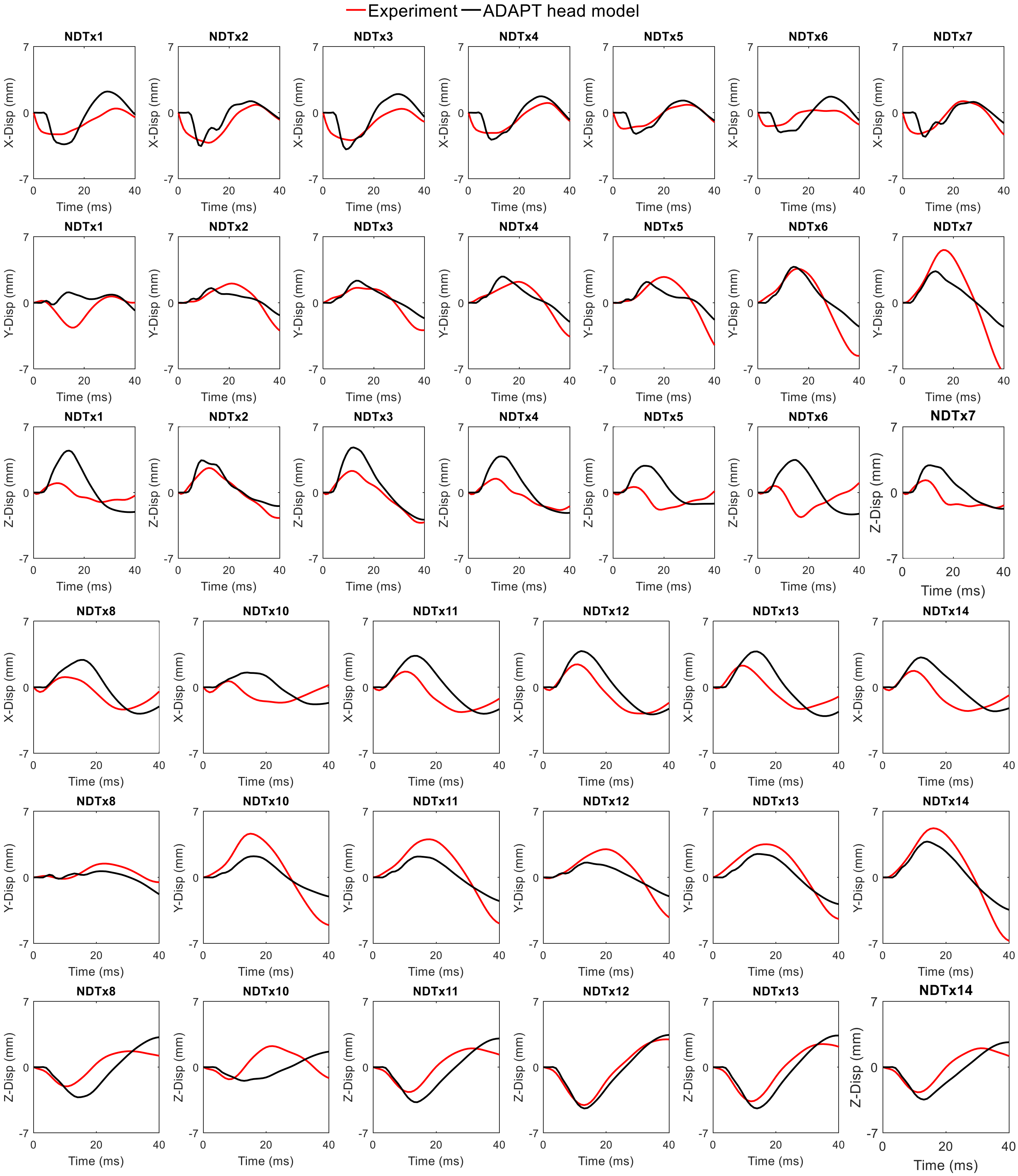


**Suppl. Fig. 2** Comparison between experimental motion and simulated brain-skull relative motion for the experiment C380-T4.

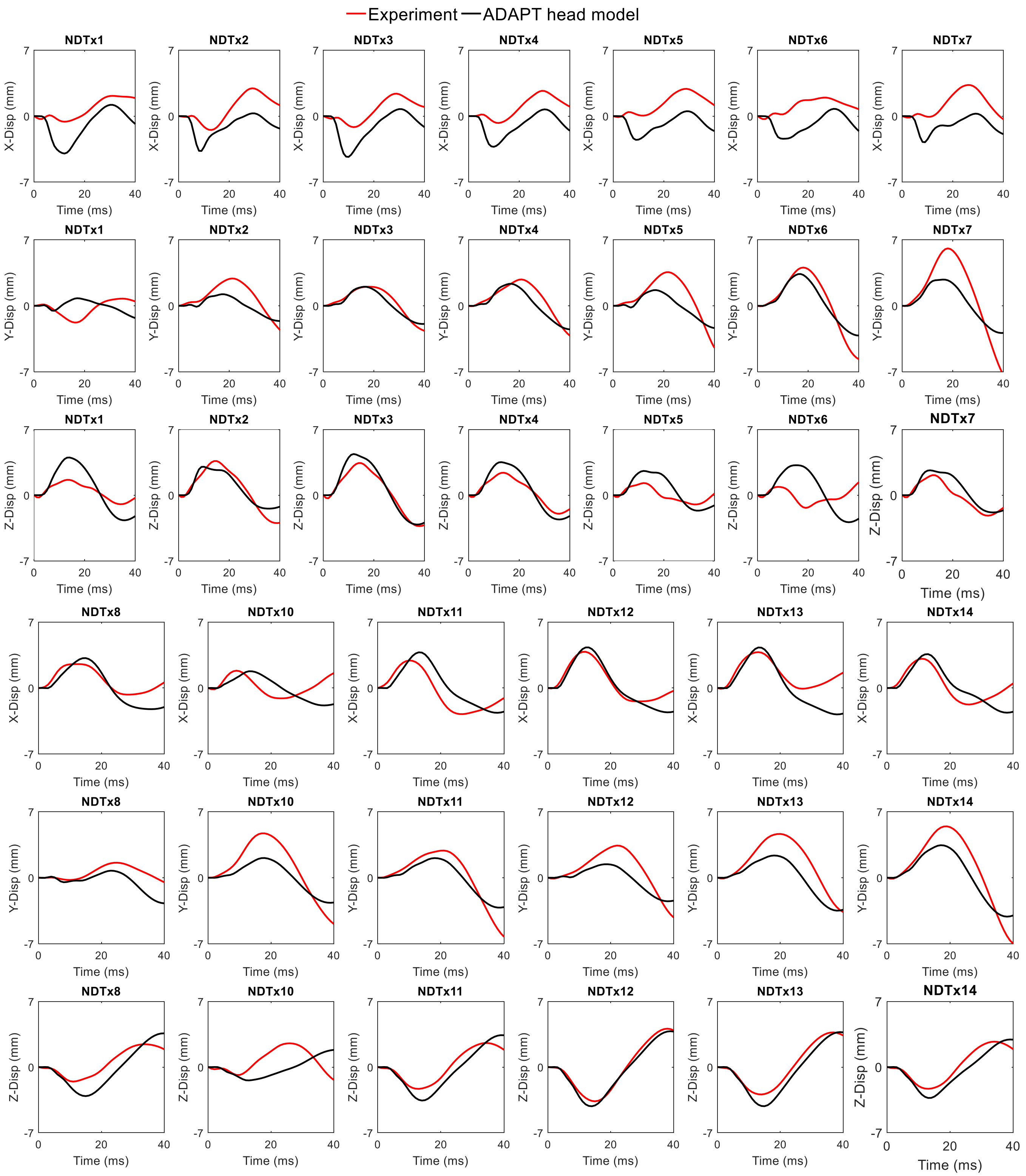


**Suppl. Fig. 3** Comparison between experimental motion and simulated brain-skull relative motion for the experiment C380-T6.

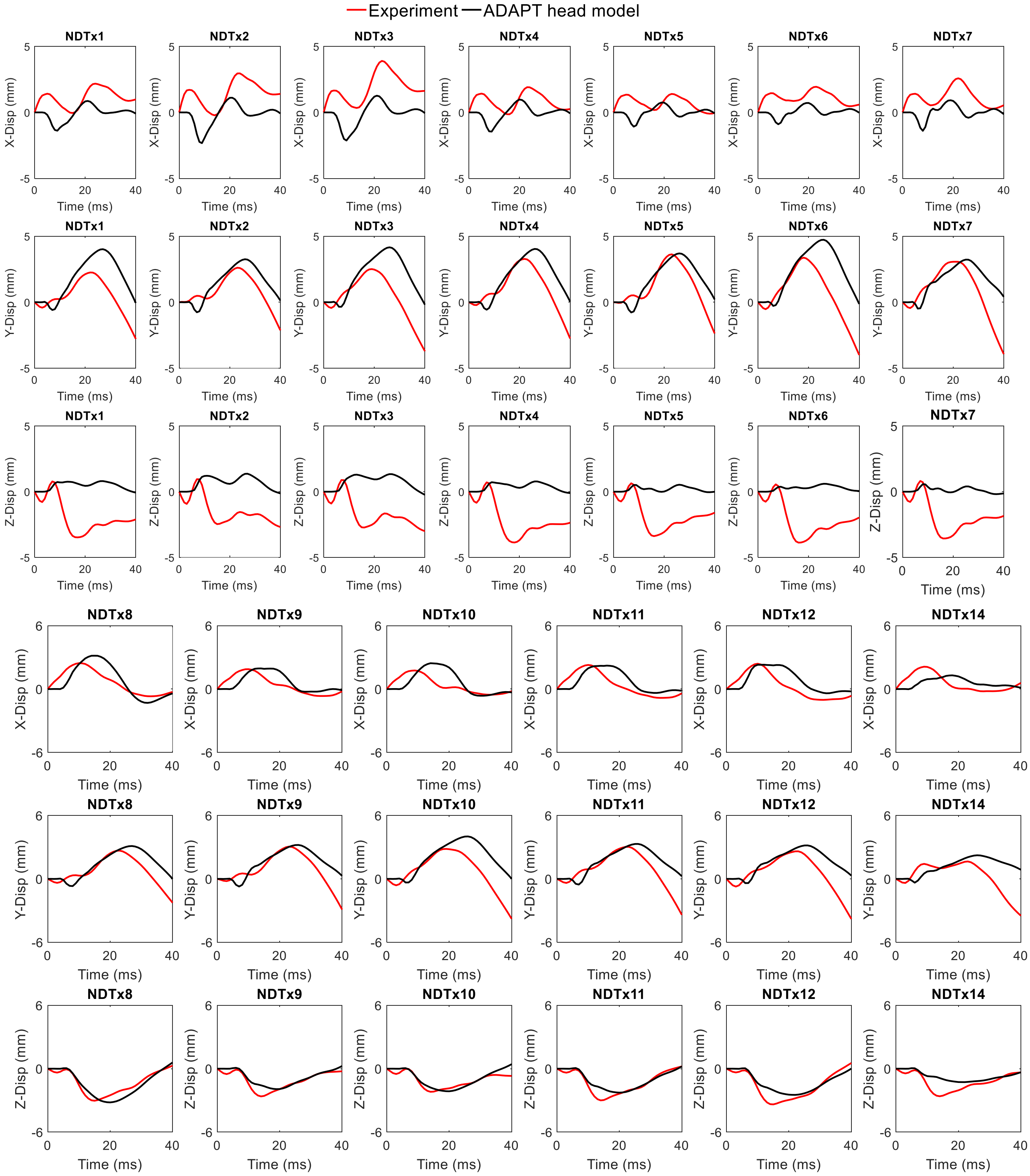


**Suppl. Fig. 4** Comparison between experimental motion and simulated brain-skull relative motion for the experiment C393-T3.

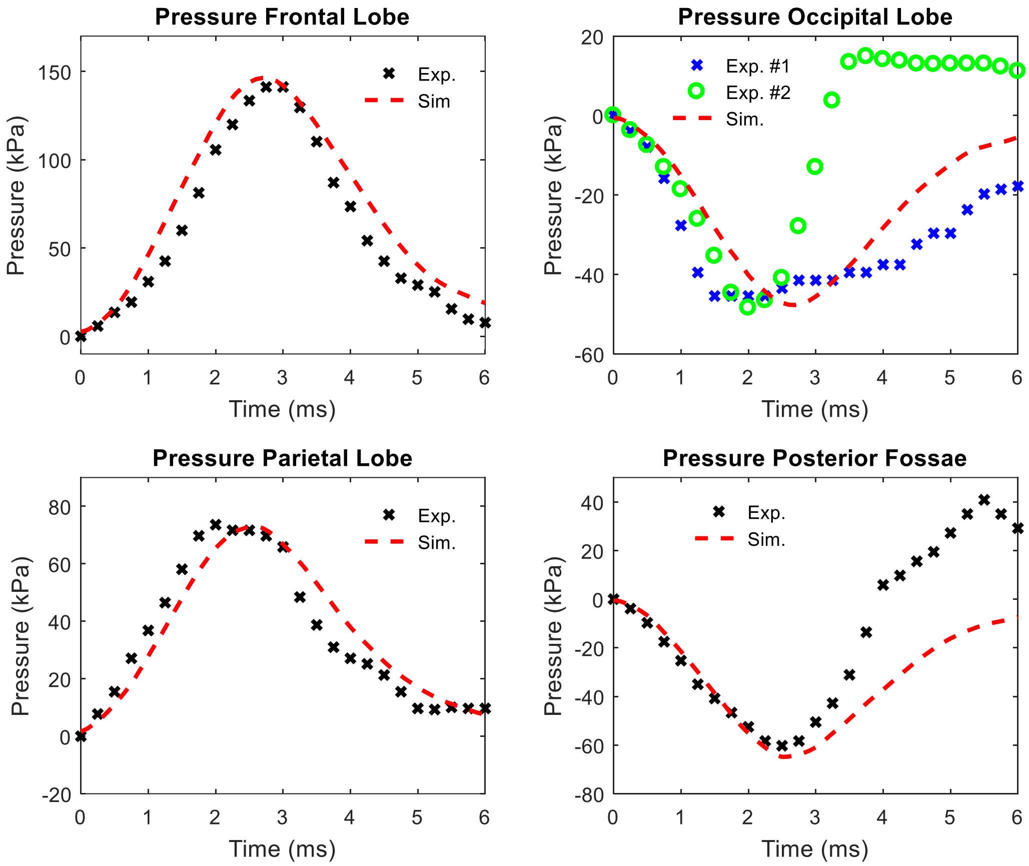


**Suppl. Fig. 5** Comparison between experimental data and simulated intracranial pressure for experiment No. 37 conducted by Nahum et al. (1977)[3].

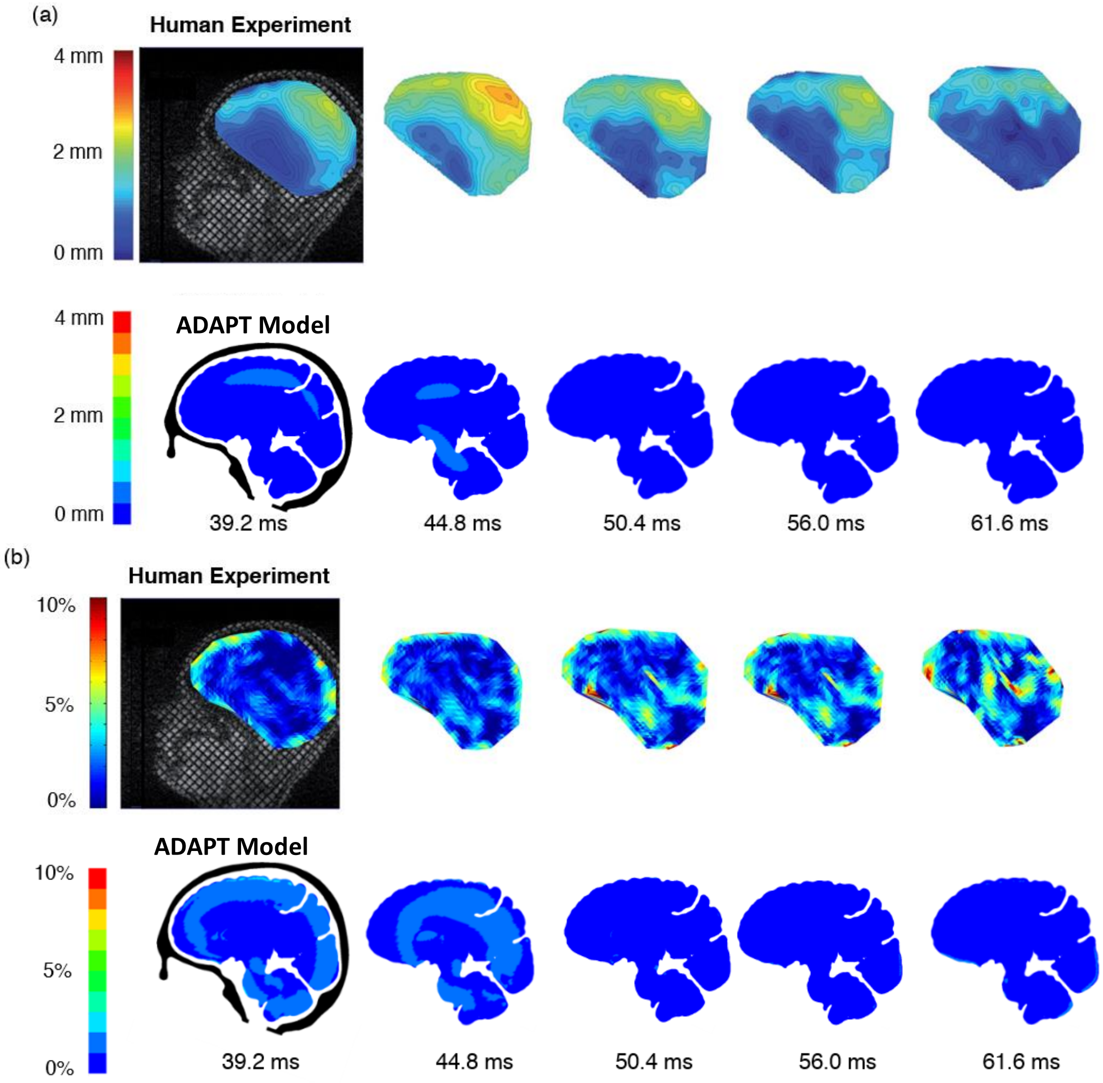


**Suppl. Fig. 6** Model prediction of skull-brain relative displacement (a) and brain strain (b) compared to a human volunteer experiment.
